# From fibril to framework: *P. abyssi* AbpX illuminates a calcium-responsive family of microbial biomatrix proteins that form thermostable hydrogels

**DOI:** 10.64898/2026.01.04.697558

**Authors:** Mike Sleutel, Adrià Sogues, Andres Gonzalez Socorro, Marcus Fislage, Vikram Alva, Han Remaut, Vincent P. Conticello

## Abstract

Evolutionary pressure on microbial communities propagating under extreme environmental conditions often results in unique structural adaptations to promote cell survival. Here, we report an investigation of AbpX, a biomatrix protein identified in cultures of the hyperthermophilic archaeon *Pyrodictium abyssi*. Under *ex vivo* and *in vitro* conditions, AbpX assembles into a para-crystalline lattice composed of semiflexible fibrils. CryoEM analysis of recombinant AbpX fibrils reveals that the precursor protein polymerizes through donor strand complementation (DSC), a process previously reported for chaperone-usher fimbriae in Gram-negative bacteria. Unlike the latter DSC protein polymers, AbpX undergoes chaperone-free polymerization in the presence of calcium ions, which are sequestered at the donor strand-acceptor groove interface between protomers in the fibril. Using a combination of cryoEM and crystallographic structural information, an atomic model is proposed for the AbpX lattice that provides insight into its potential role in biofilm formation. These findings suggest that calcium ion coordination triggers fibril assembly and pre-organizes the fibrils for incorporation into the protein lattice. Bioinformatic analysis indicates that AbpX exemplifies a distinct and broadly distributed clade of calcium ion responsive biomatrix proteins within the TasA superfamily that can be fabricated into hydrogel biomaterials *in vitro* under environmentally benign conditions.

**Significance:** Biofilms provide a protective environment for microbes that enhances resilience against environmental stressors. Secreted protein filaments constitute a major structural component of these extracellular matrices, however limited information is available on the mechanism of biofilm formation and structure of the resultant protein assemblies. Here, we report a class of biomatrix proteins that are widely distributed in bacteria and archaea. We demonstrate that one such protein, *P. abyssi* AbpX, self-assembles into fibrils and subsequently into para-crystalline lattices in response to the presence of calcium ion. We describe a mechanistic model for the structural evolution of the fibrils into an ordered protein framework that mimics the lattice structure of the *ex vivo* assembly observed during cell culture.

## INTRODUCTION

Biofilms are organized multicellular microbial communities tied together by multi-component organic matrices (1). The primary non-cellular components of biofilms are extracellular polymeric substances (EPS) that are secreted into local environments, e.g., exopolysaccharides, environmental DNA (eDNA), and fibrillar or surface-bound proteins (2). While the composition of biofilms differs between organisms, non-covalent polymers of peptides and proteins represent a major structural component of many microbial biofilms. Usually, these polymers are secreted as filaments that can self-associate into larger assemblies or interact with cellular surfaces and other polymeric substances thereby promoting biofilm formation. Structural analysis of *ex vivo* protein filaments at near-atomic resolution, in combination with computational protein structural prediction and bioinformatic mining of microbial genomic data, have led to significant advances in our understanding of the relationship between the polymeric structure of the extracellular protein filaments and biofilm formation and persistence. In addition to insight into microbial ecology, this structural information has enabled the development of synthetic biomaterials from biofilm matrix proteins, e.g., curli filaments (3) and Caf1 fimbriae (4), in which the native sequences are modified to enhance or expand function. This research has facilitated the development of engineered living materials (ELMs) (5), in which the pathogenic biofilm paradigm is inverted and, instead, medically or technologically beneficial organisms are encapsulated into synthetic protein-based biomatrices. Progress towards these applications necessitates the identification of protein polymers that can be processed into artificial biomatrices under mild conditions that minimize toxicity towards encapsulated substrates, e.g., sensitive therapeutics or living cells.

Recently, we reported the cryoEM structural characterization of cannulae (6), which are rigid protein nanotubes expressed on the extracellular surface of *Pyrodictium* species (7–10). Archaea within the family *Pyrodictiaceae* are obligate anaerobes that thrive in a hyperthermophilic (80 °C to 110 °C) deep-sea environment (11). *Pyrodictium* cells within the biofilm are connected through a network of cannulae that provides a supportive extracellular matrix through which cells can potentially exchange material to facilitate survival of the multi-cellular community in this harsh growth environment. Consequently, cannulae display high thermodynamic and kinetic stability *ex vivo* and *in vitro*, maintaining their structural integrity under physical conditions that would denature protein assemblies derived from mesophilic organisms. The structural robustness of cannulae and, more generally, biomatrix proteins from hyperthermophilic microbes has attracted scientific interest for the creation of synthetic biomaterials that retain the desirable properties of the native systems but can be rationally modified to enhance performance in non-native environments or to promote new technological applications.

During the structural analysis of cannula nanotube assemblies (6), cryoEM imaging of the *ex vivo P. abyssi* culture revealed the presence of an extracellular network composed of periodically interacting protein filaments. These narrow diameter filaments displayed a locally undulating, i.e., sinusoidal, character that promoted self-assembly into an ordered protein lattice. The morphology of these filaments was reminiscent of the recently reported structures of bacterial and archaeal bundling pili (BP) (12–15), which have been proposed to form the main proteinaceous components of the corresponding microbial biofilms. A hypothetical bundling pilin, AbpX (WP_338249486.1), was identified in a Foldseek (16) structural homology search within the proteome of *P. abyssi* AV2 using the structure of the TasA polymer (PDB: 8AUR) as a query. An AlphaFold3 (17) (AF3) prediction for the AbpX trimer revealed structural similarity to the experimental structures of bundling pili, particularly the formation of supramolecular polymers through donor strand complementation (DSC) (18), a form of non-covalent polymerization previously observed in the structures of the chaperone-usher (C-U) pili of Gram-negative bacteria as well as other bacterial and archaeal extracellular filaments (6, 12–15, 19–25). In DSC, the N-terminal donor β-strand from one protein subunit inserts into the acceptor groove of a β-sheet in a neighboring subunit to structurally complete its β-sandwich fold. AF3 structural predictions of AbpX oligomers also revealed the presence of an acidic pocket at the protomer interfaces that could serve as a calcium ion binding site and potentially mediate subunit interactions within the oligomer. As observed previously for cannulae (6), we hypothesized that coordination of calcium ions at environmentally relevant concentrations might spontaneously induce the chaperone-free polymerization of AbpX monomers into protein fibrils. However, due to extensive self-association, cryoEM reconstruction could not be successfully applied to determine the macromolecular structure of the *ex vivo* AbpX fibrils.

Here, we demonstrate that the mature AbpX protein can be produced recombinantly in *E. coli* through bacterial over-expression. Recombinant AbpX underwent facile self-assembly in the presence of calcium ions at ambient temperature to form semiflexible fibrils that displayed a similar sinusoidal periodicity to their *ex vivo* counterparts. CryoEM was employed to determine the structure of the recombinant AbpX fibrils at near-atomic resolution, which confirmed that polymerization occurred through N-terminal donor strand complementation and involved the coordination of three calcium ions at the donor strand-acceptor groove interface between sequential protomers in the fibril. Higher-order assembly occurred at increased ionic strength, which resulted in the formation of a thermally stable, self-supporting hydrogel composed of a para-crystalline lattice of AbpX fibrils. CryoEM image analysis of the AbpX hydrogels indicated that its underlying structural periodicity was related to the helical pitch of the AbpX fibril and that the lattice parameters of the network could be correlated with those previously observed for the *ex vivo* lattice. A crystal structure of sc-AbpX, a self-complemented version of the AbpX protein monomer, revealed the presence of crystal contacts that mimicked the intra- and inter-fibrillar interfaces of the AbpX fibril network, which enabled the development of a structural model for the protein lattice. Bioinformatic analysis provided evidence that AbpX-like proteins represented a compact and distinct clade in the TasA superfamily of biomatrix proteins, suggesting that the calcium ion-induced formation of para-crystalline protein networks may represent a widespread mechanism to promote biofilm formation within a diverse range of bacteria and archaea.

## RESULTS

### *P. abyssi* AbpX assembles into fibrils and fibrillar networks *ex vivo* and *in vitro*

The primary sequence of *P. abyssi* AbpX comprised 217 amino acids, of which the first 54 residues were predicted to be a signal peptidase I (SPase I) cleavage sequence (26). An additional gene sequence was located immediately downstream of the *abpX* gene oriented in the same direction, which we tentatively assigned to the same operon. This genetic locus encodes a protein annotated as a signal peptidase I (WP_338249487.1; NCBIFAM TIGR02228), which we postulate corresponded to a dedicated protease for AbpX signal sequence processing. A codon-optimized gene sequence of the AbpX protein, comprising an N-terminal methionine codon and the 163 amino acids of the mature sequence, was cloned into the high copy-number plasmid pD451-SR as a transcriptional fusion under the control of a T7 promoter (Fig. S1A). AF3 structural prediction of the mature AbpX monomer indicated that two cysteine residues were spatially proximal within the folded structure at a distance that was consistent with disulfide bond formation. Recombinant AbpX was expressed in *E. coli* strain SHuffle-T7 to promote intracellular disulfide bond formation. Expression cultures were screened for the presence of the recombinant protein under IPTG induction, and the protein was purified from the cell lysate using an experimental procedure previously optimized for recombinant expression of thermostable cannula proteins (*see Materials and Methods*) (6). Recombinant AbpX was obtained in an unoptimized yield of 16 mg of purified protein per gram of wet cell paste from *E. coli* expression cultures. The purity and identity of the recombinant AbpX protein was confirmed using SDS-PAGE analysis and electro-spray ionization (ESI) mass spectrometry (Fig. S2).

Negative stain (ns) TEM of solutions of purified AbpX in storage buffer (80 mM NaCl, 50 mM Tris, pH 7.5, 0.01 mM EDTA) indicated that the protein slowly polymerized into thin fibrils (∼2.5 nm diameter) upon incubation at ambient temperature (Fig. 1A and Fig. S3A). The 2D class averages of the fibrils (Fig. 1B) displayed a similar sinusoidal morphology to the thin AbpX fibrils observed in cryoEM images of *ex vivo P. abyssi* AV2 cultures (Fig. 1C). In contrast to recombinant AbpX, the *ex vivo* fibrils occurred primarily in the form of super-bundles in *P. abyssi* liquid cultures. The averaged power spectrum of the *ex vivo* AbpX fibrils displayed intensity on the meridian that corresponded to an axial rise of ∼46 Å (Fig. 1D), which was consistent with the inter-subunit distances between AbpX protomers in the AF3 structural predictions of donor strand complemented oligomers (6). Further 2D classification of the *ex vivo* super-bundles (Fig. 1E) indicated a para-crystalline texture in which the semiflexible fibrils (∼2.5 nm diameter) were organized into a periodic lattice based on a 2D unit cell having dimensions of 98 Å x 280 Å.

**Fig. 1.**
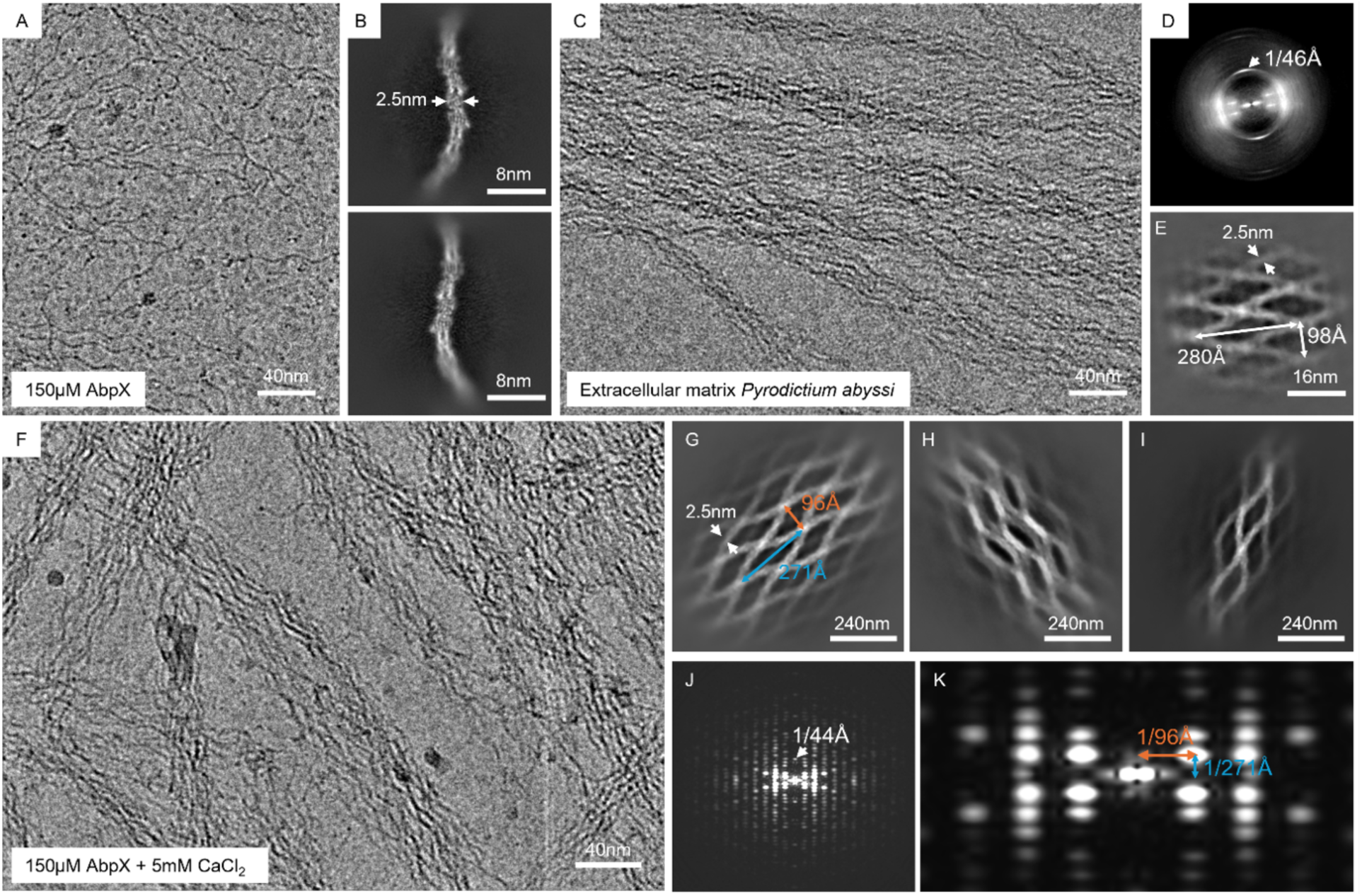
Hierarchical modes of ordering of AbpX. (**A**) CryoEM micrograph of recombinant 150 µM AbpX in the absence of CaCl_2_ shows single fibril and a background of unpolymerized single AbpX molecules; (**B**) 2D class average of AbpX fibril segments with an apparent diameter of 2.5 nm; (**C**) Para-crystalline fiber bundles found in the extracellular milieu of a *Pyrodictium abyssi* liquid culture; (**D**) Corresponding power spectrum of the *ex vivo* AbpX super bundles exhibiting a distinct maximum at 1/46 Å; (**E**) *Ex vivo* super bundle 2D class average revealing a periodic lattice of inter-connected 2.5 nm diameter fibrils spaced at 280 Å and 98 Å, respectively; (**F**) CryoEM micrograph of recombinant 150 µM AbpX supplemented with 5 mM CaCl_2_ demonstrating clustering of AbpX fibrils into laterally packed super bundles; (**G**)-(**I**) 2D class averages of recombinant AbpX super bundles, presumed to correspond to projection views of different orientations; (**J**) Power spectrum of the 2D class average shown in (**G**) revealing a distinct absence of layer lines, rather a rectangular lattice of maxima is observed. The maximum at 1/44 Å on the meridian (n = 0) corresponded to the helical rise of the fibrils; (**K**) Enlarged view of the central area of the power spectrum in (**J**) from which the typical distances in both the axial (1/271 Å) and radial direction (1/96 Å) are indicated.

To promote higher order assembly, calcium chloride (5 mM) was added to a solution of pre-formed AbpX fibrils (150 mM), which induced an immediate increase in solution viscosity that resulted in rapid hydrogelation. Negative-stain TEM (Fig. S3B-D) and cryoEM imaging (Fig. 1F) of the hydrogels indicated that the recombinant AbpX fibrils self-associated into super-bundles that mimicked the morphology of the *ex vivo* assemblies, which suggested that the fibrils were the immediate structural precursors of the lattice network. Reconstructed 2D classes corresponding to different orientational views of the recombinant bundles (Fig. 1G-I; Fig. S4 and Movie S1) revealed an underlying para-crystalline texture based on a 2D lattice having similar unit cell dimensions (96 Å x 271 Å) to that observed for the *ex vivo* bundles (6). Representative power spectra of the 2D classes (Fig. 1J-K; Fig. S4 and Movie S1) indicated a high degree of structural order within the fibril lattice. The dimensions of the 2D unit cell could be clearly discerned in the power spectrum and correlated with the spatial periodicities in the 2D classes that result from presumptive pair bonding between AbpX fibrils. The axial rise (∼44 Å) of the AbpX fibril was observed in the power spectrum (Fig. 1J) and its value closely corresponded to that observed in the power spectrum (∼46 Å) of the *ex vivo* fibrils (Fig. 1D).

### CryoEM structural analysis of recombinant AbpX fibrils

To determine the structure of the recombinant AbpX fibril, standard helical refinement procedures were employed after high-resolution cryoEM data collection on a recombinant protein sample that had not been treated with an exogenous source of calcium ions. This process afforded a 3D reconstruction of the AbpX fibril at a global resolution of 3.6 Å (Fig. S5A-G, FSC 0.143 criterion). The reconstructed density (Fig. 2A) was consistent with the presence of a fibril in which AbpX protomers were related through a left-handed 1-start helix (rise = 43.8 Å, twist = -53.97°) with a helical pitch of ∼294 Å (∼6.7 subunits/turn). The atomic model of the AbpX fibril (Fig. 2B) revealed that its structural integrity was maintained through a combination of donor strand complementation and calcium ion coordination at the structural interfaces between protomers. Despite purification of recombinant AbpX in an EDTA-containing (5 mM) buffer, the presence of coordinated calcium ions within the fibrils suggested that the AbpX protein displays a high affinity for calcium ions, which we tentatively attribute to the strong thermodynamic driving force for donor strand complementation in bundling pili and structurally related protein polymers (6, 27, 28).

**Fig. 2.**
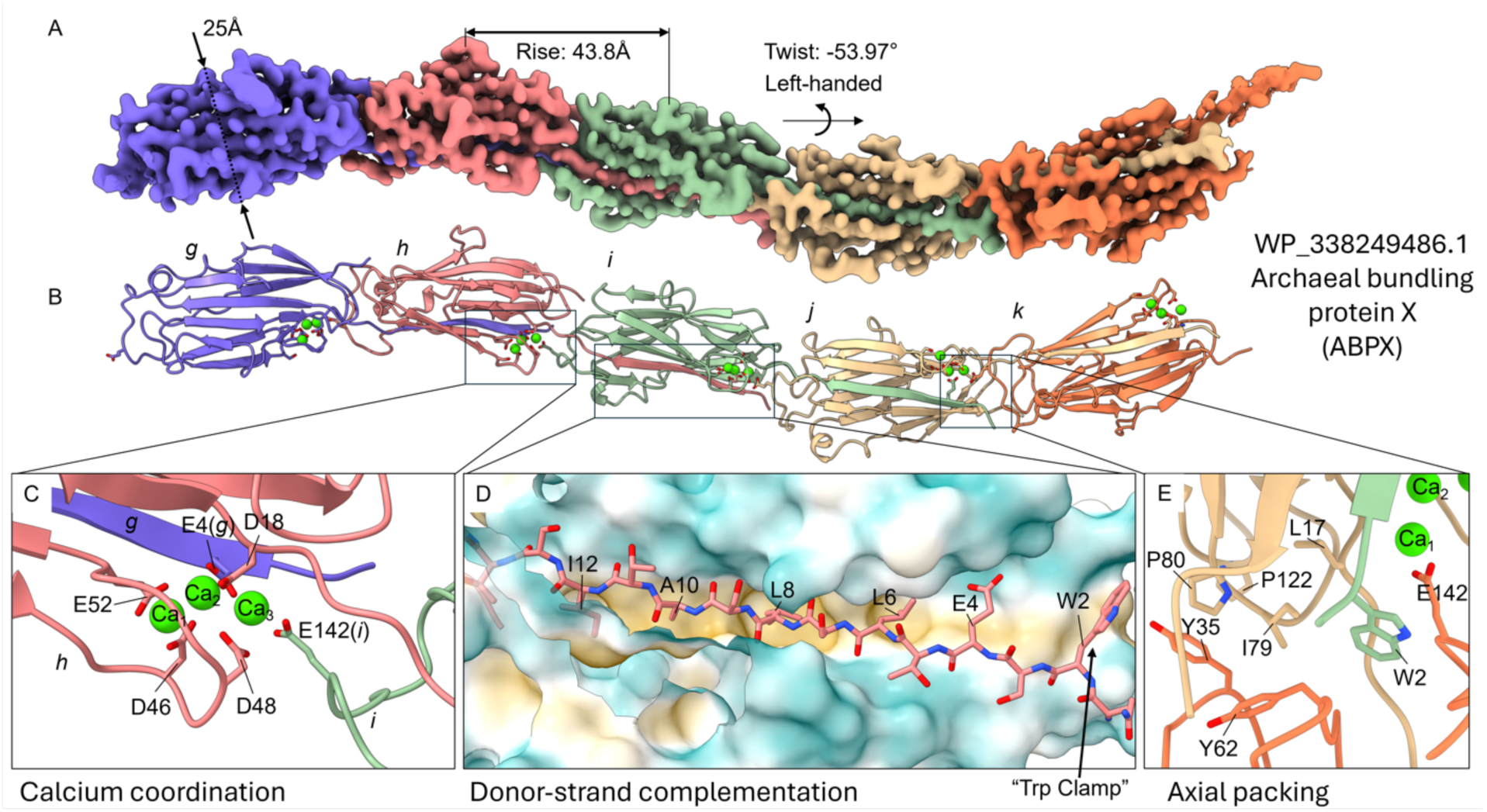
CryoEM structure of the donor-strand complemented and Ca-coordinated AbpX fibril. (**A**) Helical reconstruction of the AbpX fibril with an axial rise of 43.8 Å and a left-handed twist of -53.97° rendered at a resolution of 3.6 Å after EMReady post-processing, and colored according to molecular chains; (**B**) Corresponding molecular AbpX fibril model revealing the donor-strand complementation across neighboring protomers and the triple calcium-coordination site; (**C**) Zoom-in of the Ca^2+^-binding site of subunit *h* with the residues of three separate AbpX protomers (*h*: D18, D46, D48, E52; *g*: E4; *i*: E142) that are involved in calcium coordination shown in stick representation; (**D**) Enlarged view of the surface exposed hydrophobic groove of protomer *i* which receives the amphipathic donor-strand of preceding subunit *h* for fold complementation: strand docking is driven by the solvent burial of hydrophobic residues W2, L6, L8, A10 and I12 into complementary, hydrophobic surface depressions of subunit *i*; (**E**) Panel focusing on the axial inter-protomer interface between *j* and *k*: no hydrogens bonds were identified across this interface using the PDBePISA webserver (https://www.ebi.ac.uk/pdbe/pisa/). The interface has a predominant hydrophobic character, and we postulate that the docking of W2 serves as a “Trp clamp” that guides the donor-strand into the correct register. Additionally, E142 engages in a salt-bridge with Ca_3_, which is presumed to serve as an additional “axial rigidification” of the fibril.

Like *P. abyssi* cannulae (6), three calcium ion binding sites were observed at the structural interfaces between protomers in the AbpX fibril. However, the calcium ion binding sites in the AbpX fibril were located at the donor strand-acceptor groove interface in the fibrils (Fig. 2C) rather than at distal sites between protomers as was observed for cannulae. Acidic residues on three consecutive protomers in the AbpX fibril participated in the calcium ion binding interactions. A central protomer (*h*) contributed most of the calcium ion binding residues to the coordination sites. A preceding protomer (*g*) provided a glutamic acid residue (E4) on its N-terminal donor strand, while a succeeding protomer (*i*) added an additional glutamic acid residue (E142) that capped the third calcium ion binding site. In addition to reinforcing the donor strand-acceptor groove interaction, calcium ion coordination at inter-protomer structural interfaces potentially restricted flexibility between structurally adjacent protomers and pre-organized the semiflexible fibrils in a conformation that could potentially promote subsequent self-association into the periodic lattice of the hydrogel network (Fig. 1F). The mature AbpX protein sequence displayed a theoretical isoelectric point (pI) of ∼4.0, which reflected its compositional bias towards acidic amino acid residues (14.6% Asp/Glu). While a subset of these acidic residues defined the positions of the calcium ion binding sites (Fig. 2C), the remainder were localized on the outer surface of the AbpX fibril, contributing to its highly negative electrostatic surface potential (Fig. S5H). This high negative surface charge presumably inhibited self-association of the AbpX fibrils into a network under the low ionic strength conditions of the storage buffer. Under the high ionic strength conditions required for *P. abyssi* cell cultivation (6), AbpX was observed solely as networked fibrils (Fig. 1C). We propose that charge screening represents a plausible mechanism to control AbpX lattice formation since the high ionic strength of the marine environment would mitigate repulsive interactions between fibrils (*vide infra*).

Despite the presence of the structurally unique calcium ion binding sites, the donor strand-acceptor groove interaction in the AbpX fibrils appeared quite like other structurally characterized DSC protein polymers (6, 12–15, 19–25). AF3 structural predictions of the AbpX monomer indicated that its N-terminal extension, corresponding to the first sixteen amino acids of the mature AbpX sequence, was unstructured even in the presence of explicit calcium ions (Fig S6A). Upon donor strand complementation, the N-terminal extension adopted a β-strand conformation that completed the jelly roll fold of a succeeding protomer in the assembly (Fig. 2D; Fig. S6B). The N-terminal sequence of AbpX (W2-I12) defined an amphipathic β-strand, in which the hydrophobic residues were exposed on the strand face that packed against the acceptor groove of the succeeding protomer. The presence of an acidic residue, E4, interrupted the amphipathic character of the donor strand. However, this residue partly defined the second calcium ion binding site at the donor strand-acceptor groove interface. In the absence of calcium ion coordination, the presence of E4 could potentially inhibit premature DSC polymerization prior to protein secretion. The structural significance of this interaction was reflected in the strong conservation of an acidic residue within the donor strand of AbpX-like protein sequences (Fig. S7). In addition, we postulated that the aromatic W2 residue in the AbpX sequence performed a structurally critical role in stabilizing the donor strand-acceptor groove interactions in the fibril (Fig. 2D-E). As proposed for the structures of other DSC protein filaments (6, 19, 28–30), the W2 sidechain acted as an aromatic clamp to anchor the donor strand into its binding site; guiding the donor strand of a protomer (*i*) into the correct orientation and cross-strand register to productively interact with the acceptor groove of the succeeding protomer (*j*). In addition, the W2 residue participated in the formation of a hydrophobic cluster between two consecutive subunits (e.g., *j* and *k*) in the AbpX fibril, which provided additional axial rigidification to the structure (Fig. 2E). Multiple sequence alignments (MSAs) of AbpX-like proteins confirmed that the presence of an aromatic residue at the first position of the N-terminal donor strand was strongly conserved (Fig. S7).

The cryoEM structure of the AbpX fibril provided a useful frame of reference from which to understand the structural hierarchy of the protein lattice. Since addition of exogenous calcium ion (5 mM CaCl_2_) induced the rapid gelation of the AbpX fibrils, we posited that the structure of the hydrogel lattice must be closely related to the AbpX fibrils. The high kinetic stability of DSC protein polymers (31) and the rapid onset of gelation argued against disassembly of the AbpX fibrils prior to formation of the para-crystalline lattice. In addition, a close structural relationship between the two assemblies could be inferred from a comparison of the helical parameters of the semiflexible AbpX fibrils and the lattice parameters of the fibrillar network. The AbpX lattice was composed of thin fibrils having a sinusoidal periodicity and narrow diameter that approximated the corresponding values observed for isolated AbpX fibrils (Fig. 1B,G). In addition, the meridional spacing (44 Å) observed in the power spectrum of the recombinant AbpX lattice (Fig. 1J) closely matched the helical rise (43.8 Å) of the AbpX fibrils (Fig 2A). Despite the similarity in helical rise, the long inter-node distance (271 Å) between fibril contacts in the AbpX lattice was significantly shorter than the helical pitch of isolated AbpX fibrils (294 Å).

Careful analysis of the 2D classes derived from the fibril network revealed that the positions of the maxima in the power spectra, corresponding to the rise and pitch, 1/44 Å and 1/271 Å, respectively, of the 1-start helix (Fig. 1J,K), remained constant for AbpX fibrils embedded in the para-crystalline mesh (Movie S1). This observation supported the hypothesis that little to no structural variability/flexibility occurred across particles that were retained after 2D classification, and that the helical symmetry of the fibril was constant once it is integrated into the lattice network. This observation stood in contrast to the 2D classes of single isolated fibrils (Fig. S5C), which tended to be less well defined with blurred density away from the center of the 2D class. Moreover, the helical symmetry error surface generated by cryoSPARC during helical refinement was characterized by a “shallow” local minimum (ranging from approximately 52.3° to 55.3°). The refined helical symmetry of the consensus volume for the AbpX fibril afforded a twist value of -53.97° with respect to the vertical axis after proper assignment of the helical hand (*vide infra*), which was consistent with the presence of angular disorder within the isolated fibrils. To identify and resolve particle heterogeneity, a 3D variability analysis (3DVA) was performed on the cryoEM reconstruction of the AbpX fibril. The 3DVA calculated the principal modes of variability within the dataset of aligned particles and was used to resolve continuous variability trajectories in real-space. A principal mode was identified in the 3DVA analysis that resolved an axial flexing of the AbpX fibril (Movie S2). Taken together, these observations underscored the structural flexibility (with respect to the fibril axis and the helical twist) of isolated AbpX fibrils outside of the lattice network. The presence of cumulative angular disorder has been previously reported for protein polymers forming thin semiflexible filaments (32, 33), including those that form networked filamentous assemblies (*vide infra*).

### Crystallographic analysis of a self-complemented AbpX monomer

Despite the high degree of order in the power spectra of the bundles, cryoEM imaging did not provide a sufficiently diverse range of 2D views from which a reliable 3D reconstruction of the AbpX lattice could be performed. As an alternative approach, we employed single crystal X-ray diffraction to obtain high-resolution structural information that could provide insight into the packing of AbpX subunits in an ordered lattice. Since AbpX polymerized into fibrils over time at low concentration, we focused instead on a self-complemented version of AbpX (sc-AbpX) in which the protein sequence was circularly permuted through transposition of the N-terminal donor strand to the C-terminus of the protein through introduction of a flexible tetra-glycine loop (Fig. S1B). This approach has been employed previously to analyze donor strand-acceptor groove interactions in pilin proteins in the absence of polymerization, as well as to identify long-range non-covalent interactions (28, 30, 34, 35). Since self-complementation resulted from intramolecular binding of the donor strand to its own acceptor groove, we postulated that sc-AbpX would form soluble monomers that were resistant to intermolecular DSC polymerization. In addition, we hypothesized that self-complementation of AbpX would not inhibit calcium ion coordination or necessarily prevent the formation of other native-like structural interfaces upon crystallization.

After bacterial expression and protein purification, recombinant sc-AbpX was crystallized, and the structure was solved in space group P6_5_22. The crystal structure of sc-AbpX displayed a high degree of anisotropy in which the resolution along the c-axis, i.e., parallel to the *6_5_*-screw, reached a higher level (1.44 Å) than in the other two dimensions (2.37 Å). At the molecular level (Fig. 3A), the sc-AbpX monomer adopted a similar jelly roll fold as AbpX protomers within the fibril with the exception that the C-terminal donor strand occupied its own acceptor groove rather than that of an adjacent subunit. Matchmaker alignment of backbones of sc-AbpX monomer and the AbpX protomer (Fig. S8) afforded an RMSD of 0.562 Å over 144 out of 163 pruned atom pairs (neglecting the swapped donor strand and a conformationally mobile loop). The C-terminal donor strand of sc-AbpX folded into the acceptor groove forming a similar network of sidechain contacts to those observed in the structure of the AbpX fibril. The three calcium ion binding sites of the polymer were conserved in a similar spatial arrangement within the crystal structure of the sc-AbpX monomer. A glutamate residue (E163) in the C-terminal donor strand glutamate of sc-AbpX made an intramolecular contact to the second calcium ion that was structurally indistinguishable from the intermolecular contact of E4 in the N-terminal donor strand of the AbpX fibril.

**Fig. 3.**
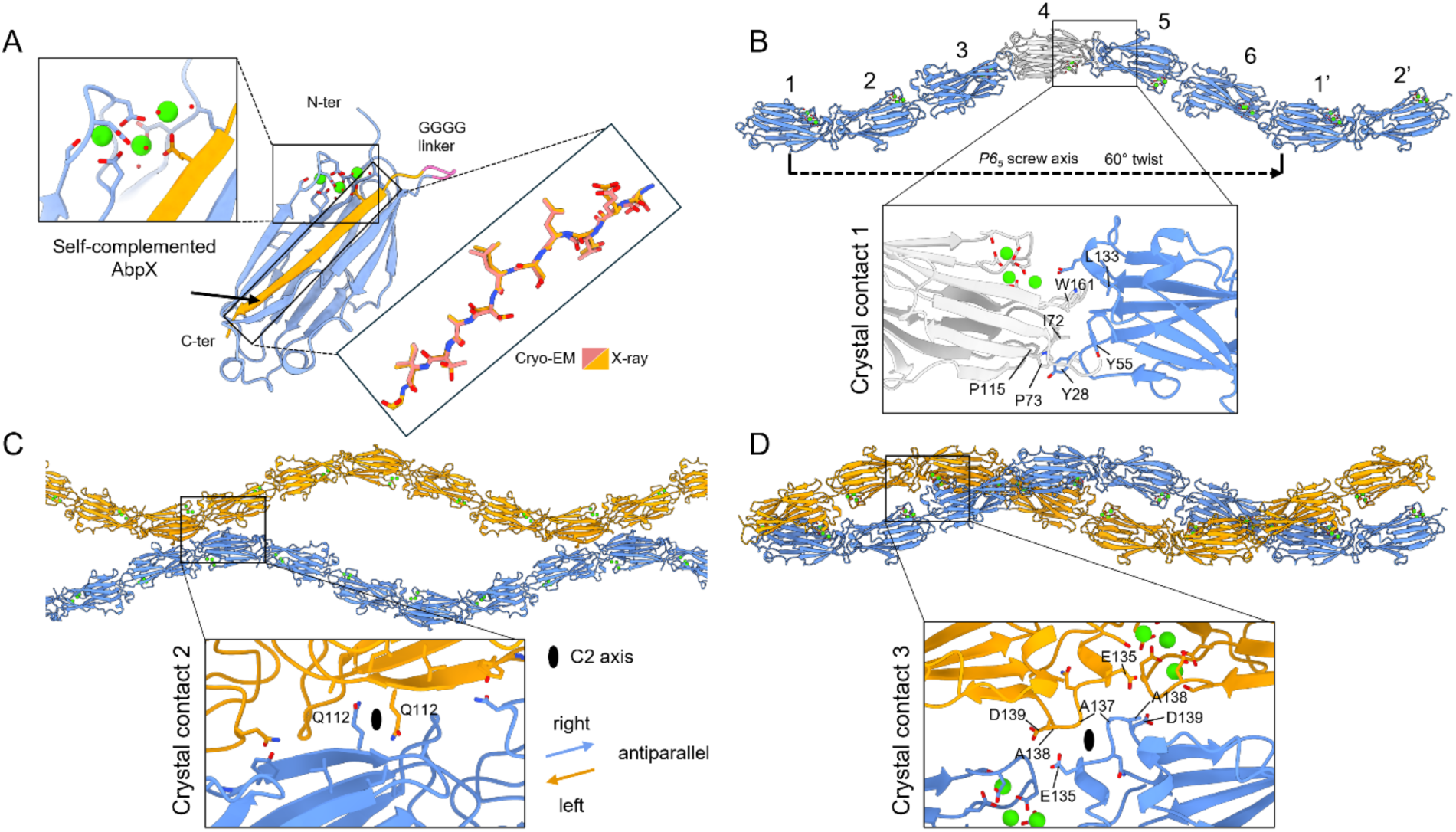
P6_5_22 X-ray crystal structure of self-complemented AbpX. (**A**) Cartoon representation of the self-complemented AbpX (sc-AbpX) monomer with the C-terminally positioned donor-strand docked into the surface exposed hydrophobic groove, and a fully occupied triple Ca-binding site. The expanded panel of the donor-strand in stick representation shows the nearly identical positioning of the donor-strand in the X-ray (yellow) and cryoEM (salmon) derived structures. (**B**) – (**D**) Crystal packing analysis of the three crystal contacts: (**B**) Crystal contact 1 (CC1): axial head-to-tail protomer packing along the P6_5_ screw axis results in a fibrillar architecture with a -60° twist and 44.77 Å rise. Inset: expanded view of CC1 that resembles the axial inter-protomer contacts present in the cryoEM model of the AbpX fibril; (**C**) Crystal contact 2 (CC2): Antiparallel fibril pair coupled via crystal contact 2 at 268.6 Å axial increments (i.e. the “fibril” pitch). Inset: expanded view of CC2 centered on the twofold axis involving 9 hydrogen bonds; (**D**) Crystal contact 3 (CC3): Second class of antiparallel fibril pairs engaged in crystal contact 3 at 44.77 Å increments (i.e. the “fibril” rise). Inset: expanded view of CC3 centered on the twofold axis of the P6_5_22 space group, involving no hydrogen bonds. C2 axes are indicated with solid lens symbols. Protofilament coloring indicates directionality: orange, left; blue, right).

Three distinct crystal contacts were observed between monomers in the crystal structure of sc-AbpX. The first crystal contact (CC1) (Fig. 3B; Fig. S9) was located along the structurally unique *6_5_* crystallographic screw axis. Despite the absence of intermolecular donor strand complementation, the sc-AbpX monomers were packed into a helical arrangement along the screw axis that was strikingly like the packing of AbpX protomers in the left-handed 1-start fibril of cryoEM structure (Fig. 2A-B). The twist (-60°) and rise (44.77 Å) of the screw axis defined a “crystallographic” sc-AbpX fibril with a helical pitch of 268.6 Å, corresponding to six subunits per left-handed helical turn. For comparison, the helical pitch length of the isolated AbpX fibril was 294 Å with 6.7 subunits per left-handed helical turn. In contrast, the long inter-node distance (271 Å), corresponding to the helical pitch of networked fibrils within the AbpX lattice (Fig. 1G), closely approximated the helical pitch of the crystallographic sc-AbpX fibril. The latter observation implied a potential structural relationship between the helical packing of AbpX fibrils in the lattice and sc-AbpX monomers at CC1 within the crystal structure. In addition, CC1 conserved the cluster of inter-molecular hydrophobic contacts observed at the interface between protomers in the AbpX fibril (Fig. 2E; Fig. S9C), which represented the primary structural interaction between successive sc-AbpX monomers along the screw axis (Fig. 3B).

CC2, the second crystal contact (Fig. 3C; Fig. S10A), derived from a series of non-covalent H-bonding interactions between two laterally adjacent, mated crystallographic fibrils of sc-AbpX. The two fibrils, related through a perpendicular C2 rotational symmetry axis, made periodic contact at a spatial frequency corresponding to the helical pitch of the *6_5_*-screw axis (268.6 Å). The 2D projection of the second crystal contact bore a striking resemblance to the 2D class averages derived from cryoEM projection images of the *ex vivo* and *in vitro* para-crystalline AbpX lattices (Fig. 1E,G). The presence of the perpendicular C2 axis required that the interacting pair of crystallographic fibrils adopted an antiparallel orientation, which implied a similar relationship between mated polar AbpX fibrils within the para-crystalline lattice (*vide infra*). These results suggested that the structural interactions between sc-AbpX monomers at CC2 were related to the interfibrillar contacts within the AbpX lattice, which, like sc-AbpX, propagated at a spatial frequency corresponding to the helical pitch of the networked fibrils (271 Å). The mating pair interface between crystallographic fibrils at CC2 involved a series of local polar sidechain/sidechain and sidechain/mainchain interactions between residues (Q27, A74, Y109, T110, Q112, L113, A114 and N154) on interacting sc-AbpX monomers related through the C2 rotational symmetry axis (Fig. 3C; Fig. S10B-C). The spatial arrangement of the backbone and sidechains of the residues involved in these interactions (except for N154) was structurally conserved in alignments of the sc-AbpX monomer and AbpX protomer (Fig. S8B-C). The latter observation argued for direct involvement of these intermolecular interactions in mediating interfacial contacts between networked AbpX fibrils in the lattice. While individually weak, the cumulative strength of these local interactions would be significantly magnified through periodic repetition along the contour length of the AbpX fibrils. Similar clusters of local polar interactions mediated lateral bundling between antiparallel oriented fibrils in the cryoEM structure of AbpA bundling pili (PDB: 7UEG) from *Pyrobaculum calidifontis* (12).

Crystal contact 3 (CC3) involved a distinct interfacial interaction between a pair of supercoiled sc-AbpX fibrils (Fig. 3D; Fig S11) that underwent periodic contact at a spatial frequency corresponding to the helical rise of the *6_5_*-screw axis (44.77 Å). As in CC2, the paired fibrils were related through a perpendicular C2 rotational symmetry axis, which necessitated an antiparallel orientation between interacting sc-AbpX fibrils. The pairing interface between fibrils at CC3 was localized on a polar loop comprising residues E135-D139 in the sc-AbpX structure. However, the electron density was not resolved beyond the β-carbon atoms for residues within this loop. In fact, to trace the main chain, the map level needed to be lowered substantially from 2.33 to 1.21. The B-factor of loop E135-D139 was also remarkably high (i.e., 110 Å^2^). These observations suggested that the polar groups of the sidechains were disordered (due to local flexibility) and were not involved in specific interactions at the interface between crystallographic sc-AbpX fibrils at CC3 (Fig. S11). A backbone alignment of the sc-AbpX monomer to the AbpX protomer indicated a significant displacement of this loop between the two structures (Fig. S8B). We hypothesized that the inter-fibrillar interactions at CC3 represented a crystallization artefact, presumably because of the applied osmotic pressure due to the presence of 14.4% (w/v) PEG 8000 in the mother liquor, and, as such, had no structural relevance to the fibril interactions within the AbpX lattice. This hypothesis appeared plausible given that AbpX lattice formation occurred at low osmotic pressure and could be triggered through an incremental increase in ionic strength.

Surprisingly, the crystallographic analysis provided negligible evidence for the presence of calcium ions at the structural interfaces between crystallographic sc-AbpX fibrils at either CC2 or CC3 despite the presence of 160 mM calcium acetate in the mother liquor. These results implied that the formation of the AbpX lattice derived from a reduction in electrostatic repulsion between the negatively charged fibril surfaces rather than specific inter-fibrillar interactions involving calcium ion coordination. Since electrostatic screening between the charged surfaces of colloidal particles depends on ionic strength (36), additional cations were assessed for the ability to induce hydrogelation of AbpX fibrils. Divalent magnesium ions (5 mM MgCl_2_) promoted hydrogelation of AbpX fibrils at similar concentrations to calcium ions, while, in contrast, higher concentrations of monovalent sodium ions (500 mM NaCl) were required to achieve the same result. TEM imaging confirmed the presence of AbpX super-bundles under both sets of conditions (Fig. S3E-H). The greater efficiency of divalent cations in inducing hydrogelation of AbpX was consistent with more effective charge screening of electrostatic repulsion between AbpX fibrils due to higher cationic charge. As observed at CC2 and CC3 of sc-AbpX, charge screening could have facilitated association between spatially local regions of reduced negative electrostatic surface potential at structurally homologous interfaces on AbpX fibrils. We postulated that charge screening between AbpX fibrils enabled the formation of a specific network of H-bonding interactions that promoted formation of the lattice and that these interfacial interactions resembled those at CC2 of sc-AbpX. In contrast, we deemed it unlikely that the non-specific association observed at the CC3 interface of sc-AbpX would be conserved in the AbpX fibril lattice.

### Construction of a structural model for AbpX hydrogel lattices

Based on evidence from cryoEM analysis of AbpX fibrils and crystallographic analysis of sc-AbpX, we propose a structural model for the AbpX para-crystalline lattice based on an ordered lattice of networked fibrils (Fig. 4A). To construct this model, the crystallographic symmetry of the sc-AbpX structure was applied after removal of symmetry mates arising from the structurally inconsequential third crystal contact CC3 (see *Materials and Methods*). The lattice model derived from layers of polar AbpX fibrils oriented in parallel that interacted with fibrils in structurally adjacent layers having the opposite polarity. Antiparallel pairs of AbpX fibrils within structurally adjacent layers are related through a perpendicular C2 axis when viewed along direction 1 in Fig. 4A. High-resolution 2D classification of cryoEM images of the AbpX lattice revealed the presence of the mating site between the paired antiparallel fibrils and its associated C2 rotational symmetry axis (Fig. 4B,C). The perpendicular C2 symmetry axis implied that the pitch of the helix, and, therefore, the pair bonding contact between mated fibrils, was based on an integral number of units per helical turn (upt). The inter-fibrillar interaction in the structural model of the AbpX lattice propagated with a spatial periodicity that approximated the helical pitch of *6_5_* screw axis of sc-AbpX (∼27 nm). The node points in the lattice model directly correlated with the mating interface observed between paired polar fibrils at CC2 in the sc-AbpX crystal structure.

**Fig. 4.**
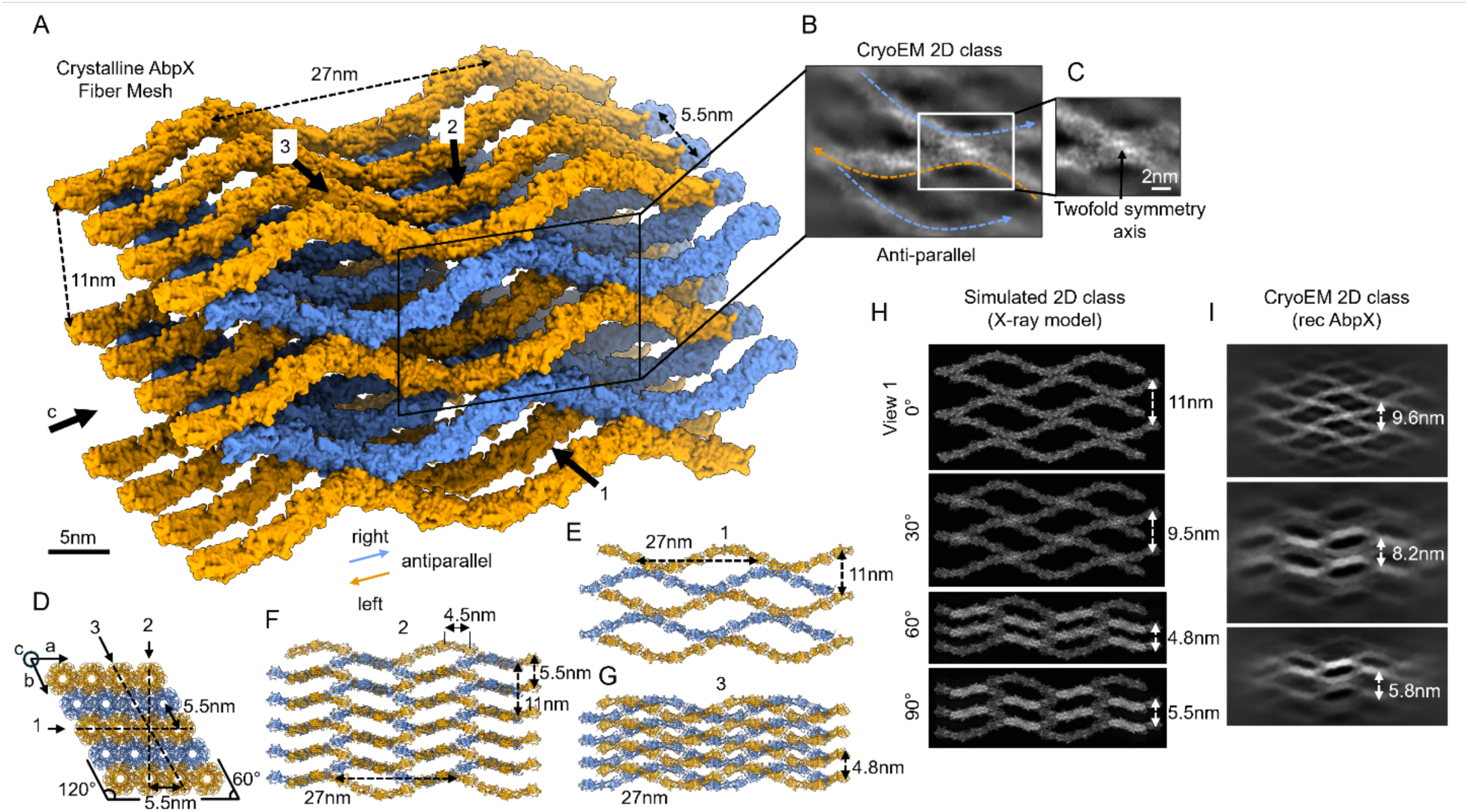
Hierarchical ultrastructure of AbpX super bundles. (**A**) Proposed model for the crystalline AbpX fibril lattice derived from the sc-AbpX structure by applying crystallographic symmetry and omitting symmetry mates that are involved in crystal contact 3. Fibrils are colored according to polarity; (**B**) High-resolution 2D class average of the antiparallel, inter-fibrillar coupling contact. Dashed, colored lines are guides for the eye mapping the directionality of the fibril; (**C**) Detailed view of the inter-fibrillar contact exhibiting twofold symmetry (in agreement with the positioning of the twofold crystallographic axis in CC2); (**D**) Cross-sectional view of the AbpX fibril lattice model: a rhombic, layered structure with constant fibril polarity along the a-direction and alternating left and right polarity along the b-direction, at 5.5 nm increments; (**E**) “Side-view”: fibril network model viewed along direction 1 (along a) shown in (**D**) with 27 nm (c-direction; pitch) and 11 nm (b-direction) periodicities; (**F**) and (**G**) Moiré patterning resulting from different viewing angles: (**F**) direction 2 (angle: 90°) and (**G**) direction 3 (angle: 60°), respectively; (**H**) Simulated 2D class averages based on the model in (a) viewed along different angles with respect to view 1; (**I**) Collection of cryoEM 2D class averages derived from recombinant AbpX super bundles, with tentative assignments of their viewing angles.

To adopt the proposed lattice model, isolated AbpX fibrils must have adjusted their helical symmetry to accommodate an integral number of protein units per turn (upt). The helical pitch of isolated AbpX fibrils was based on 6.7 upt, which argued that the most likely symmetries for the networked fibrils would correspond to 6.0 or 7.0 upt. These symmetries would require the least amount of geometrical distortion of the parent fibril (Fig. S12). Although a upt of 7 would involve only a moderate reduction in twist (from -53.97° to -51.4°), a corresponding reduction in rise of 12% (from 43.8 Å to 37.7 Å) would be required to pack into the lattice. Whereas, for the 6 upt case, the expected rise and twist were 45.2 Å and -60°, which involved only a small change in rise but a significant increase in the twist. The predicted reduction in rise for 7 upt fibril conflicted with the experimental measurement of the 44 Å rise for the crystalline mesh (Fig. S4), which was closer to the predicted value of 45.2 Å for the 6 upt fibril. Based on that observation, we tentatively concluded that the 6 upt fibrils formed the structural basis of the AbpX mesh. While the formation of the para-crystalline lattice would have required a significant increase in helical twist vis-à-vis the isolated fibrils, the angular flexibility observed in the cryoEM analysis of isolated AbpX fibrils suggested that the possibility that this geometrical distortion could be potentially accommodated through the compensatory formation of interfacial interactions between fibrils that mimicked CC2 of sc-AbpX. In addition, we postulate that charge screening due to increased ionic strength decreased the negative electrostatic surface potential on the fibril. The corresponding reduction in repulsive interactions between subunits within the AbpX fibrils could have enabled tighter winding of the helical protein fibril, potentially resulting in an increase in the helical twist. The 6 upt arrangement of subunits was observed in the “crystallographic” helix, i.e., the *6_5_*-screw axis, of sc-AbpX despite the absence of the intermolecular donor strand-acceptor groove interaction providing cohesion between protomers. The rise and pitch of the *6_5_*-screw axis, 44.7 Å and 268.2 Å, closely mimicked the corresponding values observed in the power spectra and 2D classes of the AbpX lattice, 44 Å and 271 Å, respectively.

The proposed structural model for the hydrogel lattice was based on a rhombic lattice of AbpX fibrils in which the two structurally defined interfacial contacts, CC1 and CC2, of the sc-AbpX crystal structure were retained (Fig. 4D). AbpX fibrils were aligned along the c-axis, i.e., the DSC direction, of the lattice in a manner that promoted periodic pair bonding between antiparallel oriented fibrils that were stacked in adjacent layers along the b-axis. Cut-away side views of the lattice from different directions (e.g., *a*, *b*, and *c* in Fig. 4E-G) revealed the underlying texture of the lattice from different perspectives. The projections along different directions of the lattice model were employed to simulate 2D classes (Fig. 4H), which were compared to experimental 2D classes derived from different views of the AbpX lattice observed in the cryoEM images (Fig. 4I). The remarkable degree of similarity between the simulated and experimental 2D classes argued for the validity of the proposed lattice model. However, differences were observed between the lattice dimensions of 2D classes derived from the proposed model (11 nm x 27 nm) versus the experimental projection images (9.6 nm x 27.1 nm) when viewed perpendicular to the c-axis. One confounding factor in this analysis was that the view angle could not be unambiguously identified for the experimental 2D class averages. Additionally, the sinusoidal amplitude (Fig. S13) of the AbpX filaments varied slightly between an isolated fibril and those that are embedded in the crystalline mesh, which was indicative of moderate structural variability that likely contributed to the differences between the AbpX lattice model derived from crystallographic data and the experimental 2D class averages obtained from cryoEM analysis of the AbpX meshes.

### Phylogenetic relationship of AbpX to microbial biomatrix proteins

Previous sequence-based analyses revealed that AbpX, as well as other archaeal DSC polymeric filaments, including CanA, AbpA, and the *Sulfolobus* thread subunit (Saci_0406), shared clear homology with the bacterial biomatrix protein TasA, together defining the TasA superfamily (6). A phylogenetic analysis of Tafi pili, a recently identified class of DSC protein filaments from halophilic archaea, indicated a distant relationship to AbpA and Saci_0406 (14). In support of this assertion, HHpred searches connected Tafi sequences to AbpA and Saci_0406 with probabilities greater than 90%, suggesting that Tafi should also be included within the TasA superfamily. To investigate the evolutionary origins of AbpX, its taxonomic distribution, and its relationship to other TasA superfamily members, we retrieved homologs of archaeal AbpX, AbpA, the cannula protein CanA and CanA-like proteins, Saci_0406 threads, Tafi pilin TafE, and bacterial TasA from the RefSeq database (37) using BLASTp (38). The full-length sequences of the resulting hits were pooled and analyzed using CLANS (39), which clustered proteins in a two- or three-dimensional space based on the strength of their pairwise sequence similarities estimated as from BLAST P-values. CLANS employed a force-directed layout procedure in which attractive and repulsive forces are applied according to the pairwise P-values, which caused coalescence of related sequences while weakly connected ones separated. As a result, clusters emerged for TasA superfamily proteins that reflected the underlying sequence relationships and delineated evolutionary relationships.

In the resulting cluster map (Fig. 5A), bacterial TasA homologs, exemplified by *Bacillus subtilis* TasA, formed a central, compact cluster comprising members from the phyla *Bacillota* and *Actinomycetota*. Archaeal TasA-like sequences from the phylum *Methanobacteriota* defined a distinct cluster positioned adjacent to, and tightly connected with, the bacterial TasA cluster, consistent with domain-specific diversification from a shared ancestral origin. To our knowledge, none of these archaeal TasA homologs has been experimentally characterized. Clusters corresponding to other families radiated from this central TasA cluster. AbpX-like protein sequences from the archaeal phylum *Thermoproteota* were identified as a discrete, well-defined cluster that was separated from, yet closely linked to, the central TasA cluster. Within this cluster, AbpX homologs were identified in the bacterial phyla *Bacillota* and *Chloroflexota*. Notably, archaeal and bacterial AbpX sequences did not segregate within this cluster, indicating possible horizontal gene transfer (HGT). The next most strongly connected cluster to the central TasA group were the Tafi pili (*Methanobacteriota*). The remaining families, Saci_0406 (thread), AbpA, CanA, and CanA-like, all predominantly from *Thermoproteota*, were further removed from the central clusters, indicating greater sequence divergence relative to TasA and AbpX.

**Fig. 5.**
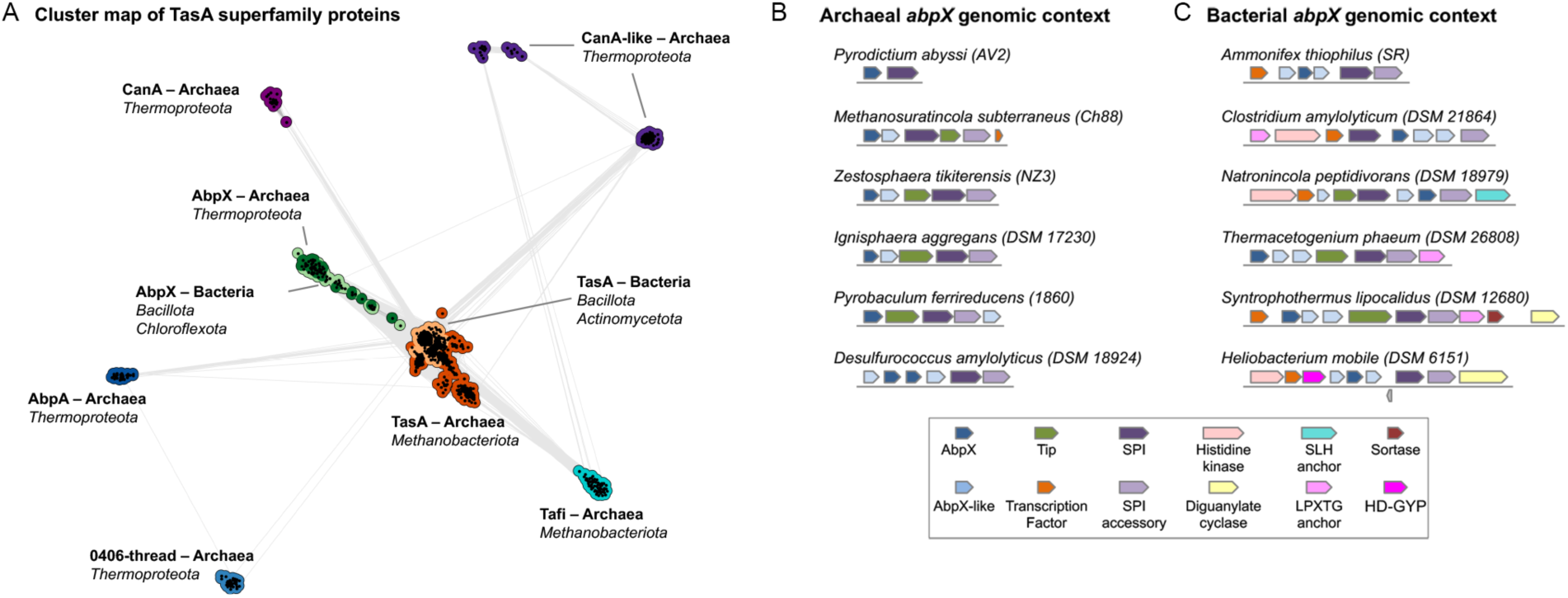
Sequence relationships and genomic context of TasA superfamily proteins. (**A**) CLANS cluster map showing the global sequence relationships within the TasA superfamily across bacteria and archaea. Each dot represents an individual protein sequence, with connecting lines indicating pairwise similarities derived from BLAST P-values; darker edges denote higher-scoring connections. Distinct clusters correspond to the major subfamilies, including AbpA, AbpX, TasA, Tafi, 0406-thread, and CanA/CanA-like groups. The analysis reveals a clear separation between the AbpX and TasA subfamilies. (**B)** Representative gene neighborhoods of archaeal *abpX* loci. In most archaeal lineages, except for *P. abyssi*, *abpX* occurs within conserved genomic contexts that encode a set of putative cell-surface-associated proteins, including SPI (signal peptidase I), SPI-accessory proteins, and predicted tip and minor filament components, together with occasional transcription factors. Although the tip components can assume different predicted structures across lineages, they are shown uniformly in green for clarity. (**C**) Representative gene neighborhoods of bacterial *abpX* loci. Unlike in archaea, bacterial *abpX* genes are frequently linked to putative surface-anchoring proteins, including SLH (S-layer homology) and LPXTG-anchor proteins, as well as sortases, and to regulatory components characteristic of biofilm-associated systems, such as histidine kinases, diguanylate cyclases, and HD-GYP (metal-dependent phosphodiesterase) signaling proteins. Across both domains, the conserved synteny patterns support the notion that AbpX proteins function within broader cell-surface modules that combine structural, regulatory, and secretion-related elements. Accession IDs for all proteins encoded by the archaeal and bacterial loci shown in panels (**B**) and (**C**) are provided in Table S3 with amino acid sequences provided in DataS1.

Like *P. abyssi* AbpX, its archaeal and bacterial homologs were characterized by low isoelectric points (pI ≈ 4.0), which reflected a high proportion of acidic residues (∼15% Asp/Glu). This compositional feature suggested that these AbpX-like proteins may also display the potential for assembly into DSC filaments in a calcium-dependent manner. Notably, AbpX homologs conserved the amphipathic N-terminal donor strand, analogous to the (W2–I12) sequence in AbpX, which included a conserved aromatic residue at position 2 and an acidic residue at position 4 (Fig. S7). CryoEM structural analysis of the AbpX fibril indicated a structurally critical role for these two residues in calcium ion mediated donor strand complementation (Fig. 2). Consistent with this hypothesis, AF3 structural predictions of representative archaeal and bacterial AbpX homologs revealed that calcium ion coordination frequently, although not universally, accompanied donor strand complementation (Figs. S14 and S15), which supported a shared structural and assembly framework with *P. abyssi* AbpX.

Like AbpX, the genes encoding AbpA, TafE, and TasA were often co-localized with a gene for signal peptidase I (SPase I). Notably, the dedicated SPase I associated with AbpX in bacteria and archaea was structurally distinct, possessing ten transmembrane (TM) helices, in contrast to the two TM helices typically observed in prokaryotic SPase I systems (40). Accessory proteins were frequently encoded alongside these fibril-associated genes. In *Bacillus*, the TasA locus included a class I signal peptidase (SipW) together with the chaperone/assembly factor TapA (41). The archaeal Tafi loci commonly included a predicted tip/adhesin, a membrane anchor, and transcriptional regulators (14). Examination of the genomic neighborhoods of AbpX homologs indicated that, unlike the minimalistic locus in *P. abyssi* genome, most *abpX*-like genetic loci occurred within larger gene clusters encoding putative accessory factors (Fig. 5B; Fig. S16), including AbpX-like minor subunits, predicted tip proteins, an SPase I, and a putative SPase I–associated accessory protein (DUF5305 family). For example, in the archaeon *Zestosphaera tikiterensis* (NZ3), the *abpX* locus encoded an AbpX-like minor fibril subunit (UniProt: A0A2R7Y7T0) and a predicted tip protein (UniProt: A0A2R7Y9R0; Figs. 5B), which AF3 predicted to co-assemble with the major AbpX pilin (UniProt: A0A2R7Y7S4) through donor strand complementation into continuous fibrils (Fig. S16A). The DUF5305 protein exhibited a modular architecture with an extracellular β-sandwich domain, two transmembrane helices, and a cytoplasmic alpha/beta fold, with homologs widespread across both archaea and bacteria. Notably, DUF5305 is sometimes fused to SPase I (e.g., UniProt E8MZY1; putative SPase I of *Anaerolinea thermophila*), supporting its functional assignment as an SPase I accessory subunit. This fusion topology, with extracellular and cytoplasmic modules flanking membrane spans, was reminiscent of the eukaryotic SPase I complex (42). Interestingly, DUF5305 is also found in Tafi loci (14, 43), suggesting that this accessory module may have played a broader role in the secretion or assembly of TasA-like matrix systems across domains.

In bacterial genomes, *abpX* gene clusters frequently encoded putative membrane anchors, for example, an S-layer homology (SLH) domain-containing cell-wall anchor in *Natronincola peptidivorans* (UniProt: A0A1I0A0H4; Fig. 5B) and an LPXTG-motif cell-wall anchor in *Clostridium amylolyticum* (UniProt: A0A1M6GRE1; Figs. 5B). AF3 predicted that the latter proteins formed DSC oligomers in combination with the AbpX fibril subunit (Fig. S16B). These neighborhoods frequently encoded two-component regulators (histidine kinases), c-di-GMP metabolic enzymes (diguanylate cyclases and HD-GYP phosphodiesterases), and Spo0A-like transcriptional regulators (Fig. 5B), which suggested that AbpX-like protein assembly and/or matrix production was coordinated with envelope anchoring and secondary messenger mediated pathways that have been broadly implicated in biofilm initiation and matrix remodeling. Taken together, these findings suggested that AbpX-like protein systems were integrated into a conserved regulatory and assembly framework paralleling TasA-based biofilm networks in bacteria. Their widespread occurrence and conserved genetic organization across archaeal and bacterial lineages indicated that TasA-like biomatrix formation represented an ancient and recurrent strategy for multicellular organization at cellular surfaces. Collectively, these findings reinforced that the TasA superfamily constituted a cross-domain clade in which common set of structural characteristics, i.e., a conserved jelly roll core, DSC-mediated polymerization, and supramolecular bundling, functionally defined distinct families of prokaryotic biomatrix proteins.

## DISCUSSION

The *in vitro* formation of lattice networks has been previously described for a structurally diverse range of protein filaments, including F-actin (44, 45), tropomyosin (46, 47), the HET-s prion domain (48), bactofilins (49, 50), and uromodulin (51–54). As observed for the AbpX fibril network, the structures of these lattices were proposed to result from periodic quasi-equivalent interactions between semiflexible protein filaments. In certain cases, specific interfacial interactions between recognition sites on the filament surface mediated lattice formation, e.g., as in the magnesium ion induced aggregation of F-actin into angle-layered aggregates and para-crystals (44, 45). Alternatively, physical changes in environmental conditions, e.g., a decrease in pH or an increase in ionic strength, induced self-association of the protein filaments into a periodic lattice through a reduction in electrostatic repulsion between negatively charged surfaces. We propose that the formation of the para-crystalline AbpX lattice was closely related to the latter situation, as no direct evidence was found for calcium ion coordination at the inter-fibrillar crystal contacts in the sc-AbpX structure. Furthermore, addition of alternative cations, e.g., Mg^2+^ and Na^+^, initiated hydrogelation of isolated AbpX fibrils, which provided further support for a mechanism in which lattice formation resulted from a reduction in repulsive interactions between protein fibrils. Increased ionic strength has been demonstrated to initiate self-assembly of uromodulin into a protein lattice (51, 52) (*vide infra*) and promote bundling of TafE pili (14) in the haloarchaeon *Natrinema* sp. J7-2. Like AbpX, the composition of the latter proteins was enriched in acidic residues, which resulted in a highly negative electrostatic surface potential on the respective protein filaments. We posited that this reduction in the electrostatic repulsion between fibrils also enabled the emergence of structurally defined pairing interactions that promoted lattice formation, as observed for CC2 in the structure of sc-ApbX and proposed for the inter-fibrillar contacts in the AbpX network (Figs. 3C and 4A).

CryoEM projection images of the *ex vivo* and *in vitro* AbpX lattices indicated that the structural parameters of the respective 2D unit cells were nearly identical (Fig. 1E,G), which implied a close structural relationship between the two assemblies despite a significant difference in assembly conditions and the potential for post-translation modification of the *ex vivo* protein fibrils. While most *in vitro* assembled protein lattices have not been associated with a corresponding biological function, it has been argued that evolutionary selection pressure is sufficiently stringent that preservation of an ordered protein lattice *in vitro* would not occur unless its formation was important for native function (55). However, a direct relationship between lattice formation and native biological function has only been established thus far for bactofilins and uromodulin. Bactofilins, a class of bacterial cytoskeletal proteins, have been observed to polymerize into filaments that further self-associate into ordered meshes *in vitro* and *in vivo* (49, 50). These protein networks have been associated with a range of biological roles including maintenance of cellular shape. However, the β-solenoid conformation of bactofilin protein promotes polymerization through formation of a cross-β fibrils (56), a comparable mode of assembly to prions (48, 57) and curli fibers (3). This polymerization mechanism distinctly differed from the DSC observed for AbpX fibrils. which suggested that the structure of the bactofilin lattice must differ significantly from the AbpX fibril network

In contrast, uromodulin, the most abundant protein in human urine, formed a similar lattice of sinusoidal fibrils through a process that we envision to be analogous to that of AbpX (53, 54). The C-terminus of uromodulin consisted of two zona pellucida domains (ZP_N_ and ZP_C_) that were separated by a flexible inter-domain linker (IDL). The ZP domains comprised structurally incomplete immunoglobulin (Ig) folds that polymerized through donor strand complementation. In the polymer, the ZP_N_ and ZP_C_ domains interacted with the IDLs on preceding and succeeding protomers, respectively, to complete the respective immunoglobulin folds. At physiologically relevant ionic strength, the resultant sinusoidal fibrils self-associated into a persistent and structurally ordered mesh that represented the physiologically functional form of uromodulin (51–53). The mannosylated N-terminal protein domains of uromodulin were presented at the surface of the polymeric lattices, where the glycans were available to bind to the tip fibrillin on type I pili of uropathogenic *E. coli*. The bacteria were entrapped within the matrix and excreted from the body through urinary shear flow. The protein lattice of uromodulin resembled the structural model that we propose for the AbpX lattice, in which DSC protein fibrils mated through formation of pairwise structural interfaces. The ability of uromodulin networks to sequester and contain microbial cells under shear flow may imply a similar functional role for the AbpX lattices in maintenance of *P. abyssi* communities under the harsh environmental conditions that the cells experience within their native ecological niche. *P. abyssi* AV2 was isolated from the porous wall of a deep-sea black smoker chimney (9), in which cells were organized into microbial communities through an extracellular matrix of protein filaments including cannulae and, presumably, AbpX fibril networks. We postulate that AbpX lattice formation may represent an evolutionary adaptation that enabled the microbial community to preserve its structural integrity under the high shear stress conditions that it would experience under the flow field of the hydrothermal vent.

Structural and bioinformatic analysis suggested that AbpX homologs formed a compact and distinct clade within the TasA superfamily of biomatrix proteins that was closely related to the canonical TasA group. Biomatrix proteins in the TasA superfamily have been directly implicated in biofilm formation (2). Structural analysis of *ex vivo* filaments within the TasA superfamily indicated that precursor proteins underwent extracellular polymerization through a similar process of donor strand complementation (15). The resultant polar filaments were anchored on the cellular surface and interacted with filaments having the opposite polarity localized on adjacent cells. These intracellular interactions promoted cohesion of the culture into a biofilm. We hypothesize that AbpX fibrils could perform a similar function to bundling pili but through a distinct mechanism in which the fibril lattices act as a scaffold that supports biofilm formation. We did not observe intact cells during cryoEM imaging of *P. abyssi* liquid cultures, and, therefore, could not gain insight into the cell-lattice interaction. In addition, while local genomic neighborhood analysis of *abpX* genetic locus indicated co-localization of a dedicated SPase I, accessory proteins were not identified within the local genomic neighborhood that could serve as a cellular anchor for the AbpX fibrils. However, accessory proteins were identified within the genetic loci of AbpX homologs (Fig 5B) that could play a similar role to that tentatively attributed to TapA (58) and AbpB (12) in maintaining cellular attachment of the TasA and AbpA filaments, respectively. Lacking definitive evidence for cell anchoring, we speculate that the unusual disk-like cellular morphology of *P. abyssi*, coupled with its preferred hyperthermophilic environment, may potentially lead to a preference for a different mode of biofilm formation, i.e., a secreted protein network rather than attached extracellular bundling cables. *Pyrodictium* species are known to produce unique extracellular structures like cannulae (7–10), which exhibit limited phylogenetic distribution and which have not been observed thus far in cultures of other cultivated archaea (6). In contrast to cannulae, putative AbpX-like protein homologs were observed to be widely distributed in archaea and bacteria, which suggested that lattice assembly from polymeric DSC protein filaments may represent a broadly employed mechanism for biofilm formation. Structural analysis of a diverse range of AbpX-like protein fibrils from organisms that proliferate under different environmental conditions should provide further insight into the relationship between lattice formation and biofilm assembly within this protein family.

In addition to lattice formation, the presence of calcium ion binding sites at the donor strand-acceptor groove interface distinguished AbpX fibrils from other structurally characterized bundling pili (12–15). We propose that calcium ion coordination serves several critical roles in AbpX assembly: (i) acting as a chemical trigger/foldase to prevent premature intra-cellular polymerization; (ii) rigidifying the fibril to promote lattice formation; (iii) inducing super bundling of the fibrils into a para-crystalline lattice; and (iv) stabilizing the protein fold as an adaptation to the extreme growth environment of *P. abyssi*. In contrast to other archaeal bundling pili, AbpX protomers did not engage in extensive axial contacts over an extended donor strand interface (Fig. S17). In the absence of calcium ion coordination, AbpX fibrils would likely be rendered too flexible to be able to self-associate into an ordered lattice. Calcium ion coordination has been previously reported to structure flexible loops and promote self-association of secreted proteins at the extracellular surface, e.g., in RTX adhesins (59) and in bacterial S-layers (60). In comparison to AbpX, previously characterized bacterial or archaeal bundling pili do not rely on calcium ion coordination to initiate donor strand complementation or to mechanically rigidify the resultant fibrils. The latter observation implies that these bundling pili may rely on alternative mechanistic pathways to facilitate donor strand complementation (58) and/or stabilize the resultant filaments, e.g., through isopeptide bond formation (Fig. S18) (13, 61, 62).

We recently reported the cryoEM structures of *ex vivo* and *in vitro* recombinant cannulae from *P. abyssi* (6), which demonstrated that calcium ion coordination reinforced and stabilized the structural interfaces between protomers even though the binding sites were not directly involved in the donor strand-acceptor groove interaction. Through a comparison between the crystal structure of a monomeric cannula protein and the cryoEM structure of a protomer in the filament, we postulated a role for calcium ion binding in remotely triggering donor strand complementation in cannulae. We hypothesized that the calcium ion gradient ([Ca]_extra_/[Ca]_intra_ ∼10^4^) between the extracellular and intracellular environment provided the driving force for self-assembly of cannulae after protein secretion at the extracellular surface. The presence of the triple calcium ion coordination site at the structural interface between three successive protomers in the AbpX fibril argued strongly for a similar process, in which calcium ion coordination induced donor strand complementation after protein signal processing and secretion. Based on our *in vitro* studies, we postulate that self-association of the nascent AbpX fibrils into a lattice would have occurred rapidly after secretion due to the high ionic strength in the marine environment. We note the remarkable convergence between these two extracellular polymerization processes in *P. abyssi*, in which calcium ion binding induces donor strand complementation, resulted in structurally divergent protein filaments that sequestered calcium ions at distinct interfaces between protomers based on similar jelly roll folds. Based on these results, we postulate that induction of protein self-assembly through calcium ion coordination may represent an evolutionary adaptation to promote rapid biofilm formation in response to an environmental trigger. Since the DSC polymerization of recombinant AbpX could be induced through *in vitro* addition of calcium ions at physiologically relevant extracellular concentrations (1-10 mM), we postulate that AbpX structural homologs represent attractive substrates for the design of synthetic protein-based hydrogel materials that can be fabricated under mild conditions that exhibit minimal toxicity towards encapsulated substrates.

## MATERIAL and METHODS

### Materials

Luria-Bertani (LB) medium was purchased from IBI Scientific (Dubuque, IA). Isopropyl β-D-1-thiogalactopyranoside (IPTG) was purchased from GoldBio (St. Louis, MO). Chicken egg white lysozyme was purchased from Research Products International (Prospect, IL). Benzonase nuclease was purchased from Merck KGaA (Darmstadt, Germany). Protease Inhibitor Cocktail Set V was purchased from Calbiochem (San Diego, CA). *E. coli* strains BL21, BL21 (DE3) and SHuffle^R^ T7 strain were obtained from New England Biolabs (Ipswich, MA). Precision Plus Protein™ Kaleidoscope™ pre-stained protein standards were purchased from Bio-Rad (Hercules, CA). Custom gene synthesis of the expression cassette for the mature AbpX sequence was performed by ATUM (Newark, CA). Carbon-coated copper grids (200 mesh) and C-flat cryoEM grids were purchased from Electron Microscopy Sciences (Hatfield, PA). Lacey carbon cryo grids and 2% (w/w) solutions of methylamine vanadate (Nano-Van) and methylamine tungstate (Nano-W) negative stains were purchased from Ted Pella (Redding, CA). All other chemical reagents were purchased from either VWR (Radnor, PA) or Sigma-Aldrich Chemical Co. (St. Louis, MO) and used without further purification.

### Expression and purification of mature AbpX

A codon-optimized gene corresponding to the expression cassette of mature AbpX (Fig. S1A) was purchased from ATUM (Newark, CA) as a sequence-verified clone in plasmid pD451-SR (Kan^R^). The synthetic gene was under control of a T7 promoter with a strong ribosome binding site in a high copy number plasmid (pUC *ori*). Purified plasmid encoding AbpX was transformed into competent cells of *E. coli* SHuffle^R^ T7 strain. Single colonies of the transformants were inoculated into Luria-Bertani (LB) medium (10 mL) supplemented with kanamycin (100 µg/mL) and glucose (1% w/v) and incubated at 37 °C for 12 h. A small aliquot of the pre-culture was inoculated into a larger expression flask containing LB medium supplemented with kanamycin (100 µg/ml) incubated in a rotary shaker at 37 °C until the O.D._600_ reached a value of ca. 0.4 absorption units (AU). Protein expression was induced by the addition of isopropyl β-D-1-thiogalactopyranoside (IPTG) to a final concentration of 1 mM and incubated at 25 °C for 4 h. Cells were pelleted by centrifugation at 8000xg for 20 min at 4 °C and exposed to multiple cycles of freeze-thaw lysis. The thawed pellet was resuspended in BugBuster^R^ Protein Extraction Reagent (5 mL per gram of wet cell) and supplemented with hen egg-white lysozyme (200 µg/mL), MgCl_2_ (2 mM), and Benzonase nuclease (250 Units). The lysis solution was incubated with shaking at 30 °C for 30 min, and the insoluble cell debris was pelleted at 10,000x*g* for 20 min. The clarified supernatant was dialyzed against a low-salt buffer (80 mM NaCl, 50 mM Tris, pH 7.5) containing EDTA (5 mM) followed by the same buffer containing 0.01 mM EDTA. The dialysate was heated to 80 °C for 20 min to denature the heat-labile proteins of the host bacterium and cooled to ambient temperature. The resultant solution was centrifuged at 14000xg for 20 min at ambient temperature. The clarified supernatant fraction was dialyzed overnight against low-salt buffer in a membrane having MWCO of 7,000 Da. The protein concentration of the dialysate was determined using BCA^TM^ Protein Assay Kit with Bovine Serum Albumin as the standard. Purified proteins achieved a concentration of 150-165 µM after purification. Protein purity was assessed using SDS-PAGE gel electrophoresis and electrospray-ionization (ESI) mass spectrometry.

### Mass spectrometry

Purified AbpX sample (1 part) was added to 4 parts of ice-cold acetonitrile (ACN) and incubated on ice for 1 h. The precipitated protein was pelleted and washed multiple times with ice-cold methanol. The final pellet was resuspended in a 50% methanol-water solution. Mass spectra were acquired on a Thermo Exactive Plus using a nanoflex source with off-line adapter. The solution of AbpX (10 mL) was deposited into a New Objective econo picotip and placed in the nanosource using the off-line adapter. A backing pressure was applied using an off-line syringe to ensure flow to the tip. A voltage of 1.0 to 2.5 kV was applied to the picotip. Typical settings employed for the analysis were a capillary temperature of 320 °C and an S-lens RF level between 30-80 with an AGC setting of 1x10^6^. The maximum injection time was set to 50 ms. Spectra were taken at 140,000-resolution at a mass/charge ratio (m/z) of 200 using Tune software and analyzed with Thermo Freestyle software.

### CryoEM imaging and structural analysis of AbpX

Quantifoil® holey Cu 400 mesh grids with 2-µm holes and 1-µm spacing were glow discharged in vacuum using plasma current of 5 mA for 1 minute (ELMO; Agar Scientific). For cryo-plunging, 3 µl of a fiber suspension was applied on freshly glow-discharged grids at 100% humidity and room temperature in a Gatan CP3 cryo-plunger. After 10 s of absorption, the grid was machine-blotted with Whatman grade 2 filter paper for 3.5 s from both sides and plunge frozen into liquid ethane at -176 °C. Grids were stored in liquid nitrogen until further use. In total, two datasets were collected: one for the recombinant AbpX fibrils (dataset 1: 7,833 movies), and one for recombinant AbpX super bundles that formed after the addition of 5 mM CaCl_2_ (dataset 2: 7,724). High-resolution cryoEM movies were recorded using SerialEM 4.1.8 on a JEOL CRYO ARM 300 microscope equipped with an omega energy filter (operated at slit width of 12 eV). The movies were captured with a K3 direct electron detector run in counting mode at a magnification of 60K with a calibrated pixel size of 0.695 Å/pixel, and a total exposure of 60 e/Å^2^ over 60 frames.

Movies were imported into cryoSPARC v4.7.0 (63) for further processing. Movies were motion-corrected using Patch Motion Correction and defocus values were determined using Patch CTF. Exposures were curated and segments were picked using the filament tracer at 10 Å separation and extracted at 4x binning. After several rounds of 2D classification, initial estimates of the helical rise and twist values were obtained based on the 2D class averages and were used in subsequent helical refinement jobs. Next, particles were re-extracted at native unbinned resolution (0.695 Å/pixel) and used as input for a second round of helical refinement, followed by local and global CTF refinement, and helical refinement. In the end, 1,629,913 fibril segments were used for reconstruction of the AbpX fibril. The resulting high-resolution volume and particle stack were used for reference-based motion correction after particle duplicate removal (1,625,594) followed by a final helical refinement. The resulting particle stack was subjected to 3D classification, and a final particle stack of 1,572,118 particles was generated after 3D class regrouping. As an initial atomic model, we used an amber-relaxed AlphaFold2 model of an AbpX trimer that was generated using ColabFold (64). The AbpX trimer was rigid-body fit into the cryoEM volume and subjected to remodeling and real-space refinement in Coot.9.8.95 (65) and phenix.refine (default settings with NCS constraints). After a single AbpX protomer was built and refined, the remaining protomers were generated by applying helical symmetry in ChimeraX. Figures were created using ChimeraX 1.9 (66). Map and model statistics are found in Table S1.

### Cloning, recombinant expression, and protein purification of sc-AbpX

The DNA sequence encoding the C-terminal self-complemented AbpX (sc-AbpX), fused to an N-terminal His-tag, was synthesized and cloned into the pASK-IBA3plus vector using Gibson assembly. The construct was transformed into *E. coli* BL21 (New England BioLabs) for expression. Plasmid sequence was verified by Sanger sequencing (Eurofins). Recombinant C-terminal self-complemented AbpX was expressed in *E. coli* BL21 (DE3) cultured in Terrific Broth (TB) supplemented with 100 µg/mL ampicillin at 37 °C. Protein expression was induced with 200 µg/L anhydrotetracycline at an OD_600_ of 0.6, followed by incubation at 25 °C overnight. Cells were harvested by centrifugation and resuspended in 100 mL of lysis buffer (50 mM HEPES pH 8.0, 300 mM NaCl, 1 mM MgCl₂, DNase I, lysozyme, and EDTA-free protease inhibitor cocktail (Roche)) at 4 °C. Lysis was performed by sonication (10 cycles of 15 seconds), and the lysate was clarified by centrifugation at 25,000 × *g* for 30 min. The supernatant was applied to a 5 mL Ni-NTA affinity column (HisTrap FF crude, Cytiva). The column was washed with 5 column volumes of Buffer A (30 mM NaCl, 10 mM HEPES pH 8.0, 10 mM imidazole) and eluted using a linear gradient of Buffer B (300 mM NaCl, 10 mM HEPES pH 8.0, 1 M imidazole). Eluted fractions were centrifuged at 20,000 × *g* for 10 min and further purified by size-exclusion chromatography using a Superdex 200 16/60 column (GE Life Sciences) equilibrated with SEC buffer (10 mM HEPES pH 8.0, 100 mM NaCl) at 4 °C. Fractions containing the target protein were pooled, concentrated, flash-frozen in liquid nitrogen, and stored at −80 °C. Protein purity was assessed by SDS-PAGE.

### Crystallization, structure determination and analysis

Crystallization trials were conducted using freshly purified sc-AbpX, concentrated to 25 mg/mL with an Amicon Ultra centrifugal filter unit (10 kDa MWCO, Millipore). Crystals were obtained via the sitting-drop vapor diffusion method at room temperature using a Mosquito nanoliter-dispensing robot (TTP Labtech, Melbourn, UK). Diamond-shaped crystals of AbpX appeared after 10 days in condition E11 of the JCSG-plus screen (Molecular Dimensions), which contains 0.16 M calcium acetate hydrate, 0.08 M sodium cacodylate pH 6.5, 14.4% (w/v) PEG 8000, and 20% (v/v) glycerol. Crystals were harvested using nylon loops and flash-cooled in liquid nitrogen. X-ray diffraction data were collected at 100 K on beamline Proxima 1 at the SOLEIL synchrotron (Gif-sur-Yvette, France). The data was shown to be anisotropic, and we processed it using AutoProc and Staraniso (67) to a 1.44 Å in one direction, whereas in the other directions it reached 2.37 Å. The crystal was integrated in the P 6_5_ 2 2 space group. The crystal structure was determined by molecular replacement using Phaser (68) from the PHENIX suite (69) and using the AbpX monomer, as predicted by AlphaFold3, as the search model (70). Iterative model building and refinement cycles were performed using COOT (65), phenix.refine (71) and BUSTER (72) to reach final R values of R_work_/R_free_ of 0.203/0.249. The crystallographic statistics are shown in Table S2. Atomic coordinates and structure factors have been deposited in the protein data bank (PDB) under the accession code 9RHP. Proteins, interfaces, surfaces, and assemblies (PISA) analysis (73) of the crystal contact interfaces was performed using the PDBePISA webserver (https://www.ebi.ac.uk/pdbe/pisa/).

### Fiber crystal mesh modelling

The model for the crystalline AbpX fiber mesh was produced by applying crystallographic symmetry operations to the sc-AbpX structure (9RHP) starting model. In brief, a single sc-AbpX monomer was loaded into ChimeraX v1.9 (66), and 3x3x3 (a,b,c) crystallographic unit cells were generated using the “unitcell” command. The Unit Cell command reads in the symmetry information from the header of the input pdb file (i.e. spacegroup: P65 2 2; unit cell dimensions: 55.453 Å, 55.453 Å, 268.594 Å; unit cell angles: 90°, 90°, 120°). The rotational and translational symmetry, as defined by entry 179 of the International Tables for Crystallography (https://onlinelibrary.wiley.com/iucr/itc/Ac/ch2o3v0001/sgtable2o3o179/) (74), was employed to populate the unit cells. Symmetry mates produced by crystal contact 3 (Fig. 3D) were manually removed, leading to the proposed model for the fibril crystal mesh shown in Fig. 4.

### Bioinformatics

To retrieve sequences for cluster analysis, we queried the RefSeq databases (37) using BLAST (38) for homologs of representative TasA superfamily members, including *Bacillus subtilis* TasA (UniProt P54507), *Pyrodictium abyssi* AbpX (NCBI WP_338249486.1), *Pyrobaculum calidifontis* AbpA (UniProt A3MUL8), *Sulfolobus acidocaldarius* Saci_0406 (UniProt Q4JBK8), CanA (UniParc UPI0024181436), and CanA-like proteins from *Metallosphaera yellowstonensis* (NCBI WP_009071828.1, WP_009072785.1, WP_009072754.1, and WP_009071324.1). Full-length sequences of all BLAST hits were extracted, and signal peptides were predicted using DeepTMHMM (75) and SignalP 6.0 (76); only sequences predicted to contain a signal peptide by at least one method were retained. We further filtered the dataset to retain only sequences ranging from 100 to 350 amino acid residues in length, consistent with the typical size range of characterized TasA superfamily members. Redundancy was reduced with MMseqs2 (77), retaining sequences with a maximum pairwise identity of 95% and a minimum alignment coverage of 70%. This yielded 1,931 (6) sequences, which were clustered with CLANS (39, 78) based on all-against-all BLAST P-values (cutoff 0.1) and iterated to equilibrium in a 2D space under default parameters. Genome neighborhood analysis was performed using the EFI Genome Neighborhood Tool (EFI-GNT) with default parameters (79).

## SI Supplement

The PDF file includes Supplementary Text, Figs. S1 to S18, Tables S1-S3, Legends for Movies S1 and S2, and Legend for Data S1. Other Supplementary Material for this manuscript includes the following: Movies S1 and S2 and Data S1.

## ACKNOWLEDGMENTS

High-resolution cryoEM imaging of AbpX fibrils and lattices was performed at the VIB-VUB Facility for Bio Electron Cryogenic Microscopy (BECM). We thank Dirk Reiter for assistance in data collection. This study was supported by the Robert P. Apkarian Integrated Electron Microscopy Core (RRID: SCR_023537) at Emory University, which is subsidized by the School of Medicine and Emory College of Arts and Sciences. M.S. and V.P.C. thank Ed Egelman for helpful discussions

## Author contributions

Conceptualization: M.S. and V.P.C. Investigation: M.S., A.S., A.G.S., M.F., and V.A. Visualization: M.S., A.S., A.G.S. and V.A., Supervision: M.S., H.R., V.P.C. Writing–original draft: M.S., V.A. V.P.C. Writing–review & editing: M.S., A.S., A.G.S., M.F., V.A., H.R., and V.P.C.

## Data and materials availability

The cryoEM reconstruction map for AbpX was deposited in the Electron Microscopy Data Bank with accession numbers of EMD-54935. The fibril model was deposited in the Protein Data Bank with accession number 9SJ2 (pdb_00009sj2). Atomic coordinates and structure factors have been deposited in the protein data bank (PDB) under the accession code 9RHP (pdb_00009rhp). All other data needed to evaluate the conclusions in the paper are present in the paper and/or the SI Supplement.

