## Supplementary Information for "From fibril to framework: *P. abyssi* AbpX illuminates a calcium-responsive family of microbial biomatrix proteins that form thermostable hydrogels"


**This PDF file includes:**

Supplementary Text
Figs. S1 to S18
Tables S1 to S3
Legends for Movies S1 to S2
Legend for Data S1

**Other Supplementary Materials for this manuscript include the following:**

Movies S1 to S2
Data S1

**A**

>AbpX
MKRETATPGGAPSGGSGAGGPAAGDARKTRMKPLVLLVALLPLLALTAIGLTYAQWSETLSLSATISTGS
LDVDIKANEATDVTTSSQYIDVSLSLTSTDSDQGNEQLSITIEKAYPGAEVNVTFTLENVGTIPALATIKLNT
DTIPSDVAACVNVKLYNAQGNPINTPYTIQLAPGEFEKFKLGIEIPSSCDLEENKKDAIQLNNIVEVDVEQNV
S*

ATGCAATGGTCTGAAACCCTGTCTCTGAGCGCAACGATTAGCACCGGTTCCCTGGACGTCGATATCA
AGGCAAATGAAGCAACCGATGTTACTACGAGCAGCCAGTACATTGACGTTAGCCTGAGCCTGACCTC
GACTGATAGCGACCAGGGCAACGAACAACTGAGCATTACGATTGAGAAAGCCTATCCGGGTGCTGA
AGTCAATGTCACCTTCACCTTGGAAAACGTTGGTACCATTCCGGCGCTGGCGACGATCAAGCTGAAT
ACGGACACGATCCCGAGCGATGTGGCTGCGTGTGTGAACGTCAAGTTGTACAACGCGCAGGGCAAC
CCGATCAATACCCCGTATACCATCCAGCTGGCGCCTGGCGAGTTTGAGAAATTCAAACTGGGTATCG
AGATCCCAAGCTCCTGCGACCTGGAAGAGAACAAAAAAGACGCCATTCAACTGAATAACATTGTTGA
AGTTGATGTGGAGCAAAATGTGAGCTAATAA

**B**

>sc-AbpX
MHHHHHHGSLDVDIKANEATDVTTSSQYIDVSLSLTSTDSDQGNEQLSITIEKAYPGAEVNVTFTLENVGTI
PALATIKLNTDTIPSDVAACVNVKLYNAQGNPINTPYTIQLAPGEFEKFKLGIEIPSSCDLEENKKDAIQLNNI
VEVDVEQNVGGGGQWSETLSLSATISTG*

**Fig. S1. Amino acid sequences and gene sequences for the recombinant proteins analyzed in this study.** (A) Full amino acid sequence and codon-optimized gene sequence of *P. abyssi* AV2 AbpX (WP_338249486.1). Recombinant mature AbpX, corresponding to the putative mature, i.e., signal-sequence processed, protein, was expressed as a transcriptional fusion downstream of an initiation codon (AUG) under the control of a phage T7 promoter. The maximum likelihood positions for secretion signal peptidase cleavage were determined using the PrediSi webserver (http://www.predisi.de/index.html) (*26*). (**B**) Full amino acid sequence of self-complemented AbpX (sc-AbpX).

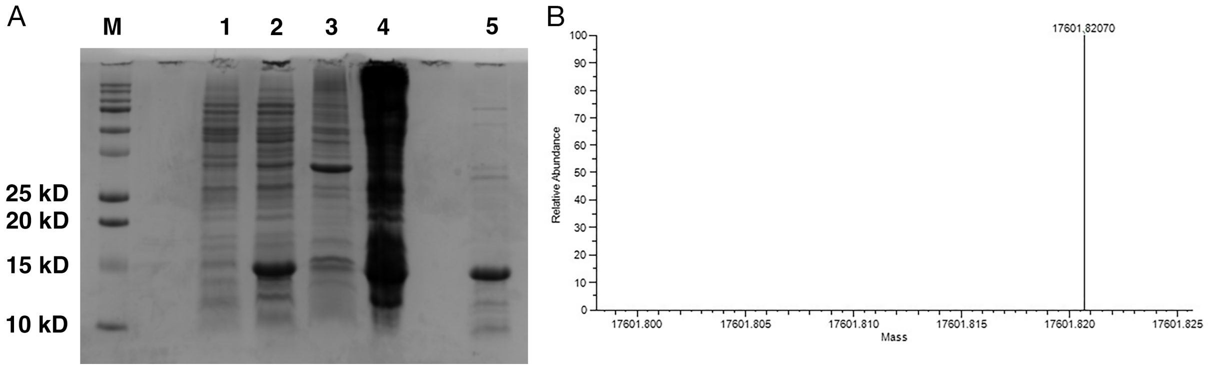


**Fig. S2. Expression and purification of recombinant AbpX**. (**A**) SDS-PAGE analysis of bacterial expression cultures and purification of recombinant AbpX. Lanes: M, protein molecular mass standards; 1, pre-induction bacterial cell culture; 2, post-induction (4 h) bacterial cell culture; 3, crude cell lysate; 4, heat-treated cell pellet; 5, purified fraction containing soluble cannula-mimetic proteins. (**B**) Deconvoluted electrospray-ionization mass spectrum (ESI-MS) of AbpX. The single peak corresponds to the disulfide-bonded mature AbpX protein (calculated monoisotopic mass: 17,602.76 Dalton, experimental monoisotopic molecular mass: 17,601.82 Dalton).

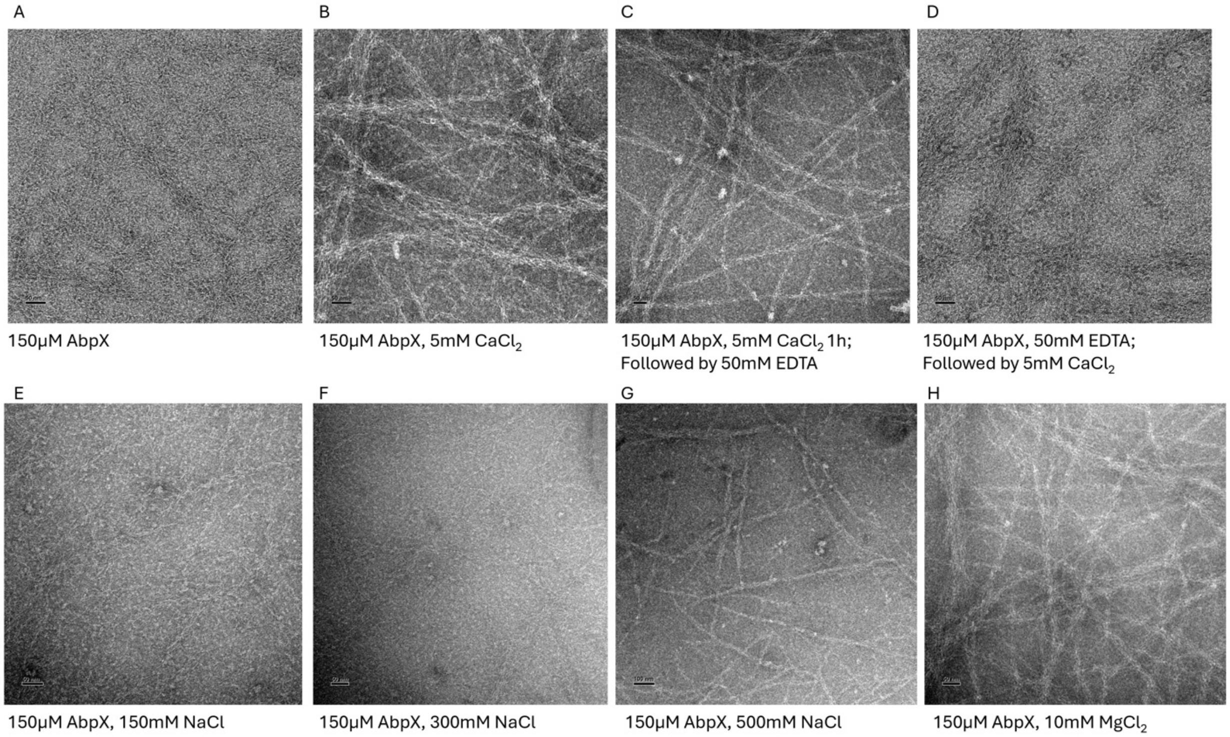


**Fig. S3. Representative TEM images of AbpX fibrils under different preparative conditions.**

.

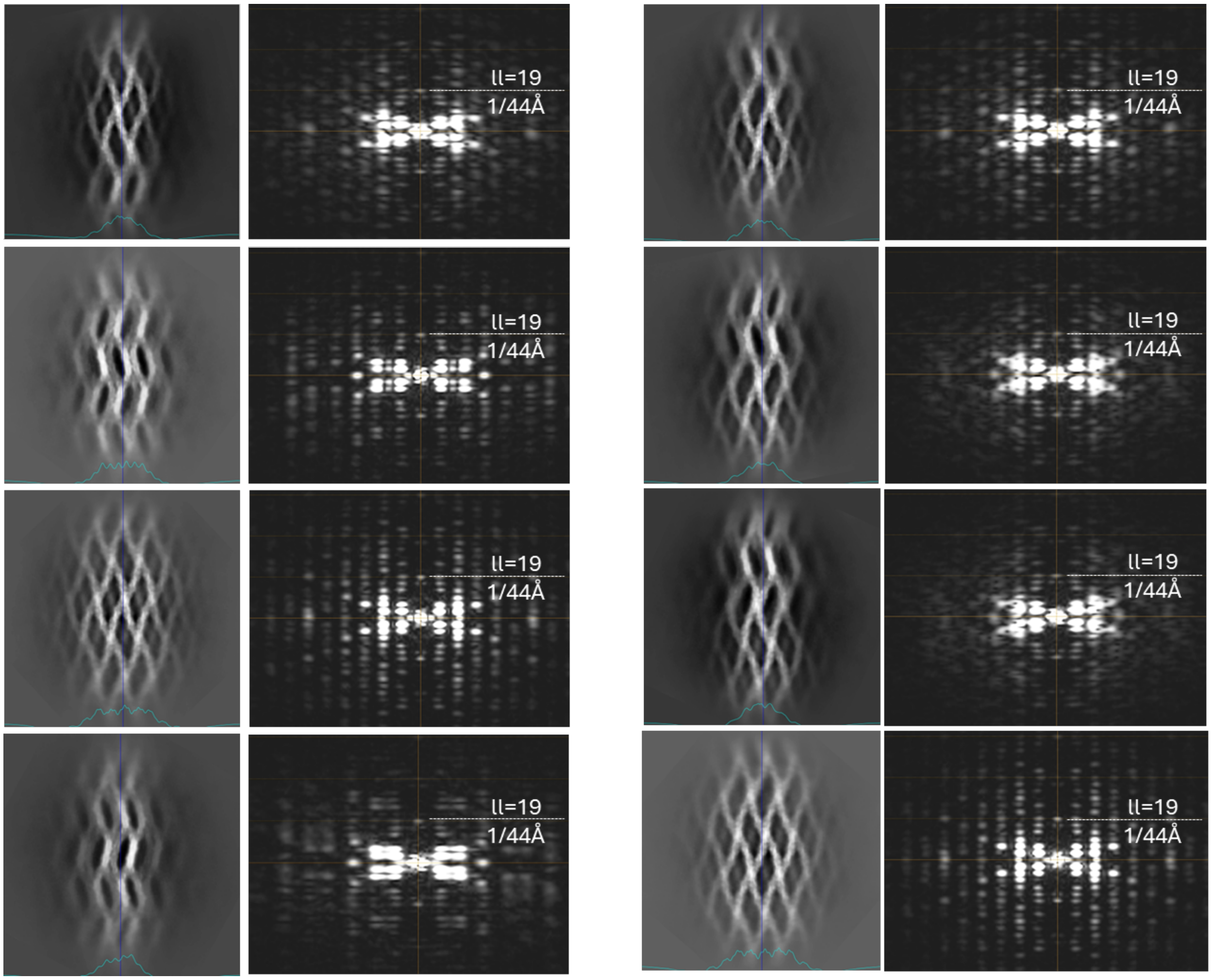


**Fig. S4. CryoEM analysis of semi-crystalline AbpX fiber bundles.** Selected frames of Movie S1 reporting 2D class averages and corresponding power spectra of recombinant, *in vitro* assembled AbpX mesh structures.

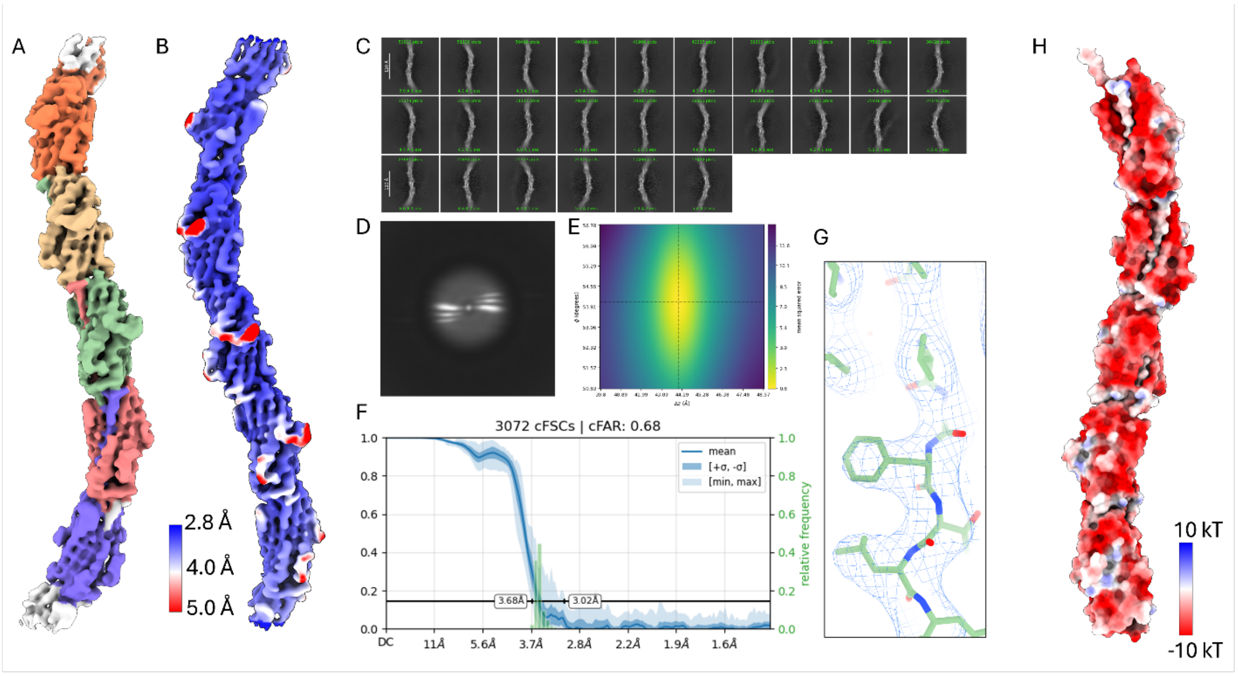


**Fig. S5. Helical reconstruction of the AbpX fibril.** (**A**) Reconstructed cryoEM volume with one fibril color-coded according to chain identity. (**B**) Surface coloring according to local resolution. Map generated by Local Filtering in cryoSPARC. (**C**) 2D class averages: vertical box size measures 278x278Å. (**D**) Averaged power spectrum of AbpX fibrils. (**E**) Helical symmetry error surface with a minimum at an axial rise of 43.8 Å and a left-handed twist of -53.97°. (**F**) The global resolution estimated by the map:map conical Fourier Shell Correlation (FSC) curve produced by Orientation Diagnostics job in cryoSPARC (v4.6.2); (**G**) Illustrated segment is used to represent the map quality. (**H**) Surface coloring according to electrostatic potential; Images were created using ChimeraX v 1.9.

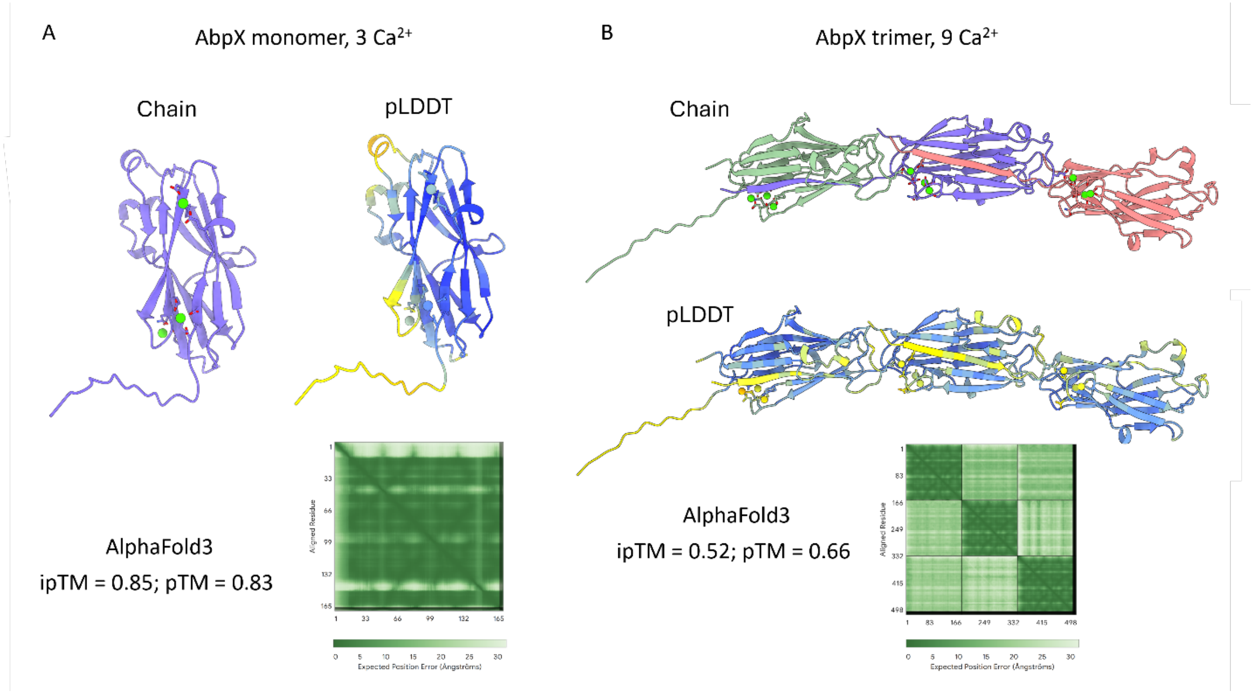


**Fig. S6. AlphaFold3 predictions of an AbpX monomer and AbpX trimer in complex with $Ca^{2+}$ ions.** Reliability metrics of the structural prediction are represented through color-coding of the model based on pLDDT and numerical values for pTM and ipTM scores. Predicted aligned error (PAE) for the AF3 structural predictions are provided as separate plots for the monomer and trimer.

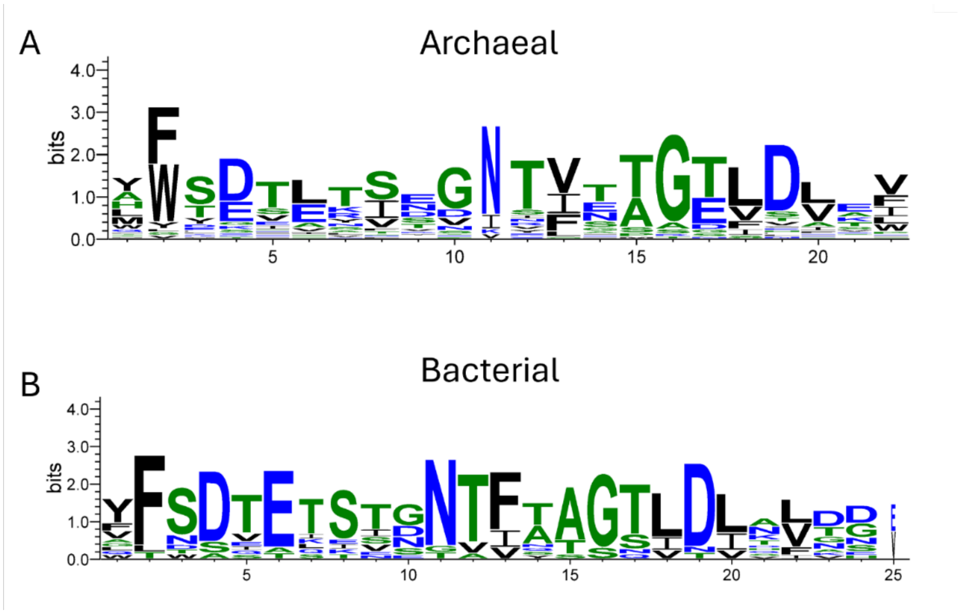


**Fig. S7. Consensus logos of the archaeal (A) and bacterial (B) donor strand sequences.**

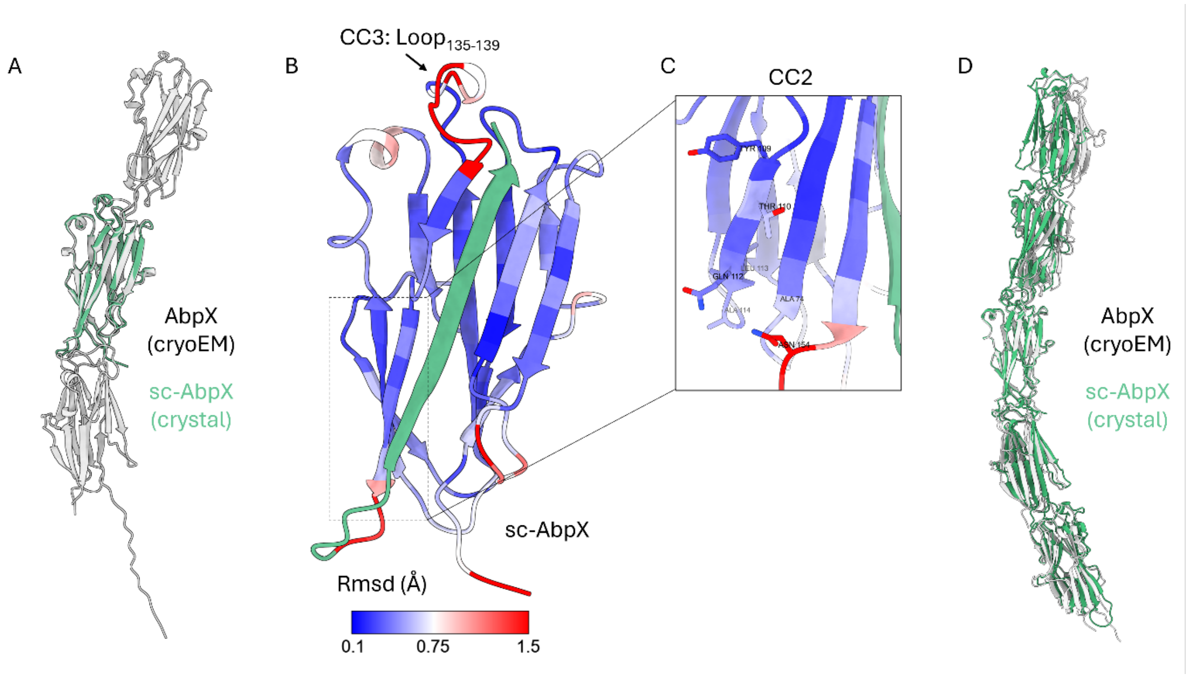


**Fig. S8. Structural comparison between AbpX and sc-AbpX.** (**A**) Structural overlay between AbpX (grey) and sc-AbpX (green). (**B**) sc-AbpX monomer colored according to root mean-squared displacement (RMSD) with respect to AbpX after a matchmaker alignment in ChimeraX 1.9. The tetra-glycine linker and the donor-strand (green) were omitted form the backbone alignment due to inherent structural mismatch that results from circular permutation of the protein sequence. Loop residues 135-139 constitute lattice contact 3. (**C**) Structural conservation of the residues involved in crystal contact 2. (**D**) Structural overlay between AbpX (grey) and sc-AbpX (green) fibrils.

Sc-AbpX: Crystal contact 1

A

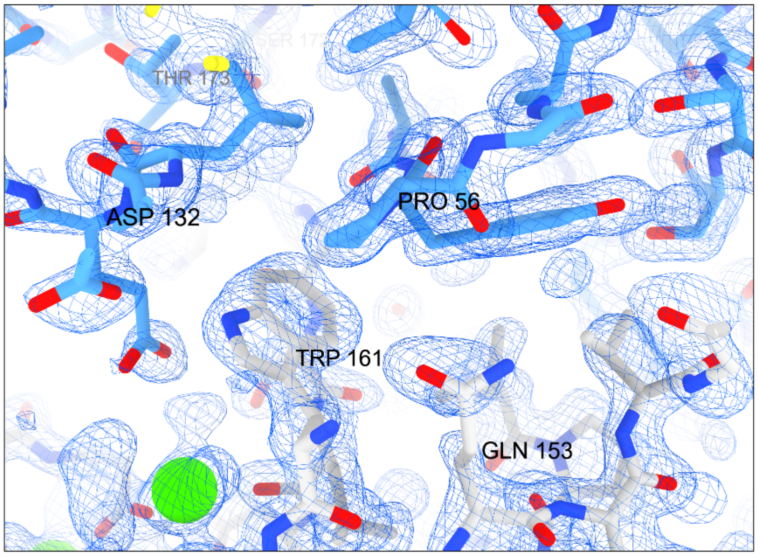


B

H-bonds

| Structure 1 | Dist. [Å] | Structure 2 |
|---|---|---|
| A:HIS 6[ N ] | 3.78 | A:SER 172[ OG ] |
| A:GLN 153[ NE2] | 3.83 | A:PRO 56[ O ] |
| A:TRP 161[ NE1] | 3.29 | A:ASP 132[ OD1] |
| A:HIS 6[ O ] | 2.60 | A:LYS 53[ N ] |

Salt bridges

| Structure 1 | Dist. [Å] | Structure 2 |
|---|---|---|
| A:HIS 6[ N ] | 4.00 | A:GLU 52[ OE2] |
| A:HIS 7[ ND1] | 3.84 | A:THR 173[ O ] |

C

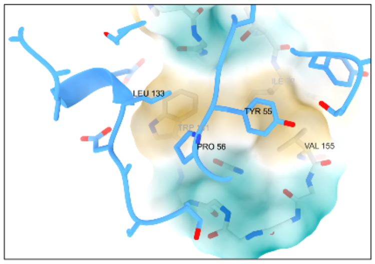


**Fig. S9. Crystal contact analysis of sc-AbpX.** (**A**) Hydrogen-bonding network of crystal contact 1 (CC1) of sc-AbpX in space group P65 2 2. Map rendered at level 1.27. Water residues hidden for clarity. (**B**) PISA analysis of crystal contact 1 (*73*). (**C**) Cartoon and surface representation of CC1 colored according to molecular lipophilicity potential (i.e. hydrophobicity).

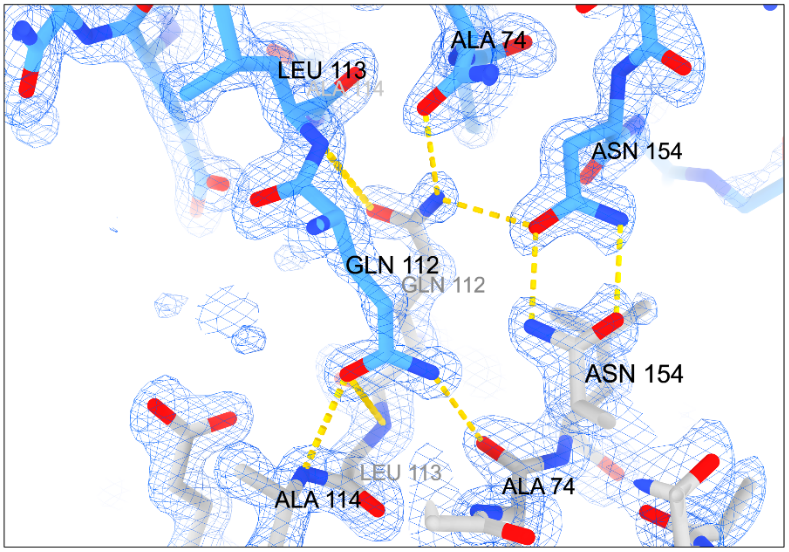


B

H-bonds

| Structure 1 | Dist. [Å] | Structure 2 |
|---|---|---|
| A:GLN 112[ NE2] | 3.36 | C:ALA 74[ O ] |
| A:GLN 112[ NE2] | 3.01 | C:ASN 154[ OD1] |
| A:LEU 113[ N ] | 3.65 | C:GLN 112[ OE1] |
| A:ALA 114[ N ] | 3.08 | C:GLN 112[ OE1] |
| A:ASN 154[ ND2] | 2.81 | C:ASN 154[ OD1] |
| A:ALA 74[ O ] | 3.36 | C:GLN 112[ NE2] |
| A:GLN 112[ OE1] | 3.65 | C:LEU 113[ N ] |
| A:GLN 112[ OE1] | 3.08 | C:ALA 114[ N ] |
| A:ASN 154[ OD1] | 2.98 | C:ASN 154[ ND2] |

C

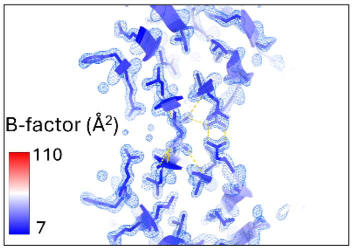


**Fig. S10. Crystal contact analysis of sc-AbpX.** (**A**) Hydrogen-bonding network of crystal contact 2 (CC2) of sc-AbpX in space group P65 2 2. Map rendered at level 2.33. Water residues hidden for clarity. (**B**) PISA analysis of crystal contact 2 (*73*) (**C**) Cartoon and stick representation of CC2 colored according to B-factor ($Å^2$).

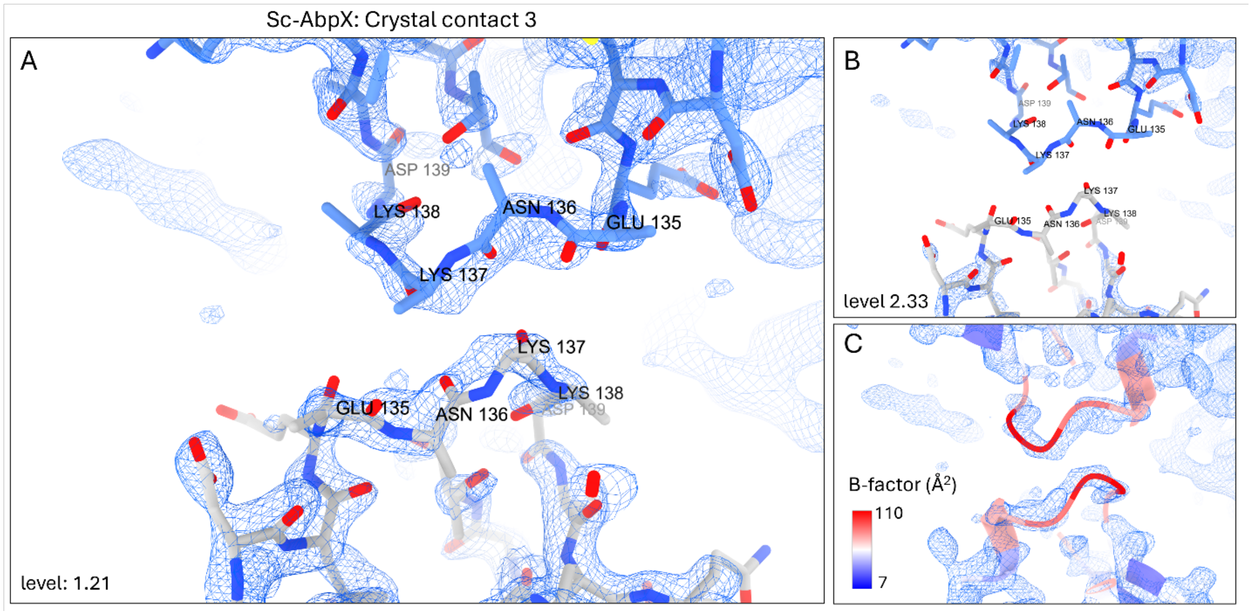


**Fig. S11. Crystal contact analysis of sc-AbpX.** (**A**) Crystallographic interface of crystal contact 3 (CC3) of sc-AbpX in space group P65 2 2. Water residues are hidden for clarity. Map rendered at level 1.21. (**B**) Map rendered at level 2.33 (i.e. the same value as used in Fig. S10A) indicated a lack of clear continuous density for the main chain; (**C**) Cartoon representation of CC3 colored according to B-factor (Å$^2$). A PISA analysis of CC3 did not identify any H-bonds and/or salt bridges.

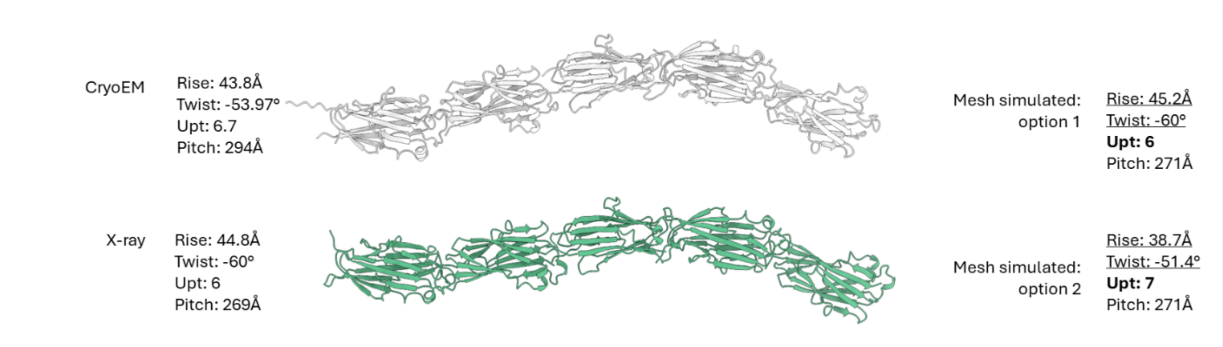


**Fig. S12. Comparative analysis of the helical parameters of the AbpX fibril.** Summary of the differences in helical rise, twist, units per turn (upt) and resulting pitch derived from the cryoEM AbpX fibril structure, the sc-AbpX X-ray crystal structure, and the simulated AbpX mesh. For the latter, we propose two options, with either 6 or 7 upt and calculate the helical symmetry that would result in either case. Crucially, for both mesh options, the pitch is derived from an experimental measurement, whereas we impose a fixed upt (shown in bold) which follows from the requirement that the upt corresponds to an integral number of protomers for the mesh. The reported rise and twist are calculated based on the pitch and imposed upt (calculated, i.e., non-experimentally determined, values are underlined) assuming a fixed sinusoidal amplitude.

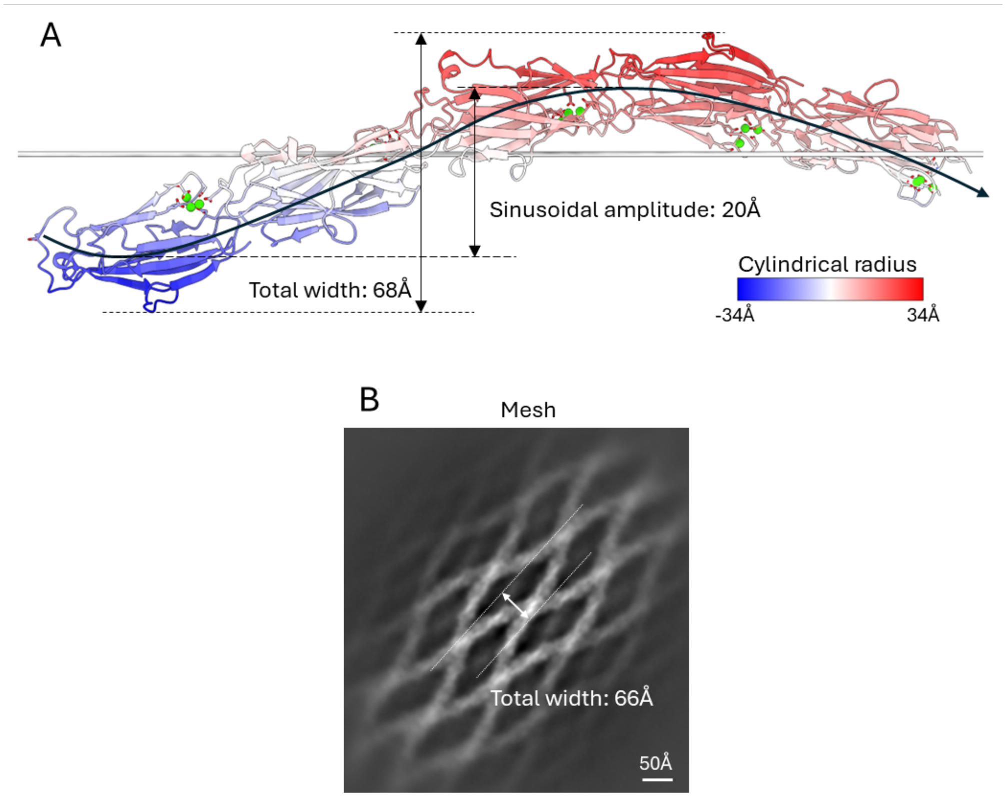


**Fig. S13. Sinusoidal amplitude of AbpX fibrils.** (**A**) Illustration and measurement of the sinusoidal amplitude and total fibril width of a solitary AbpX fibril. The structure is colored according to the cylindrical radial distance from the fibril axis. (**B**) Measurement of the total width of a mesh embedded AbpX fibril based on a cryoEM 2D class average.

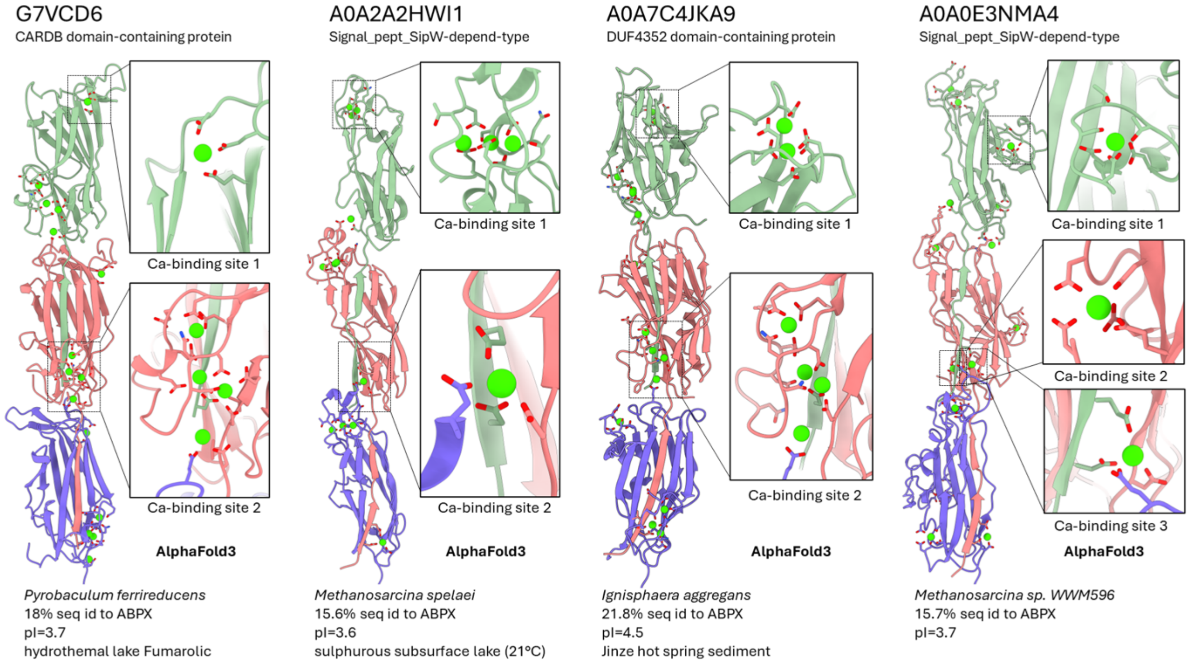


**Fig. S14. AlphaFold3 predictions of archaeal AbpX-like, Ca-coordinated DSC architectures.** Selected AF3 predictions of archaeal AbpX-like filaments with (multiple) different acidic (Asp, Glu) clusters, predicted to coordinate (multiple) calcium ions.

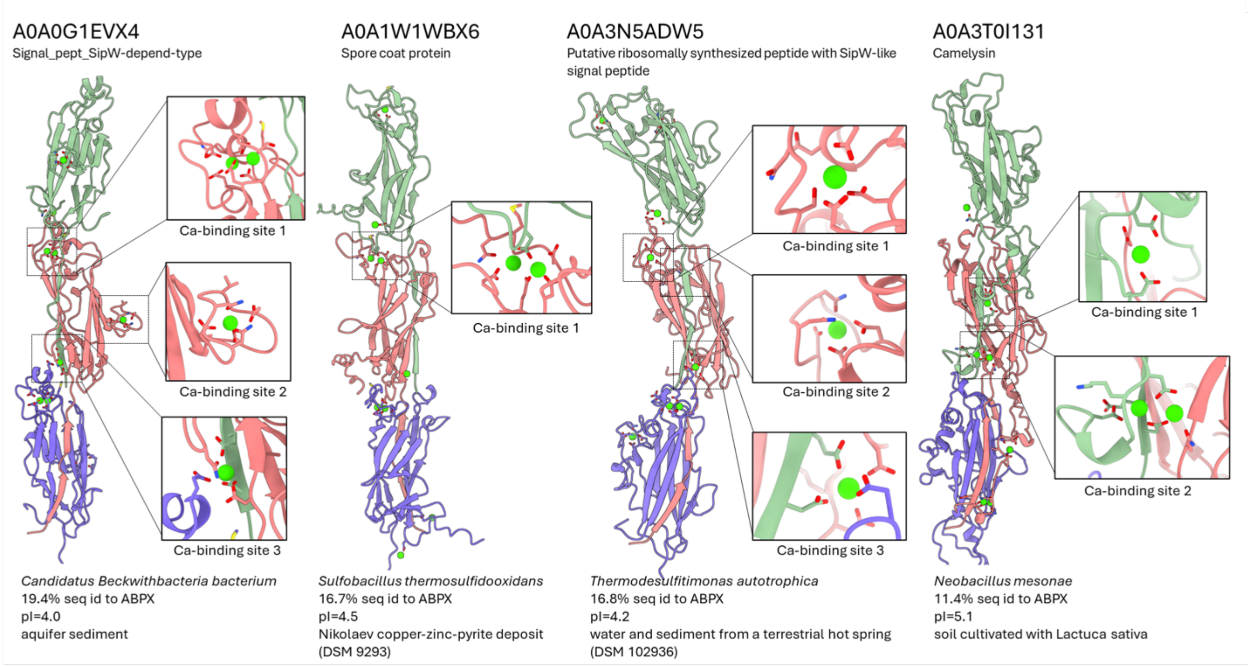


**Fig. S15. AlphaFold3 predictions of bacterial, AbpX-like, Ca-coordinated DSC architectures of AbpX homologues.** Selected AF3 predictions of bacterial AbpX-like filaments with (multiple) different acidic (Asp, Glu) clusters, predicted to coordinate (multiple) calcium ions.

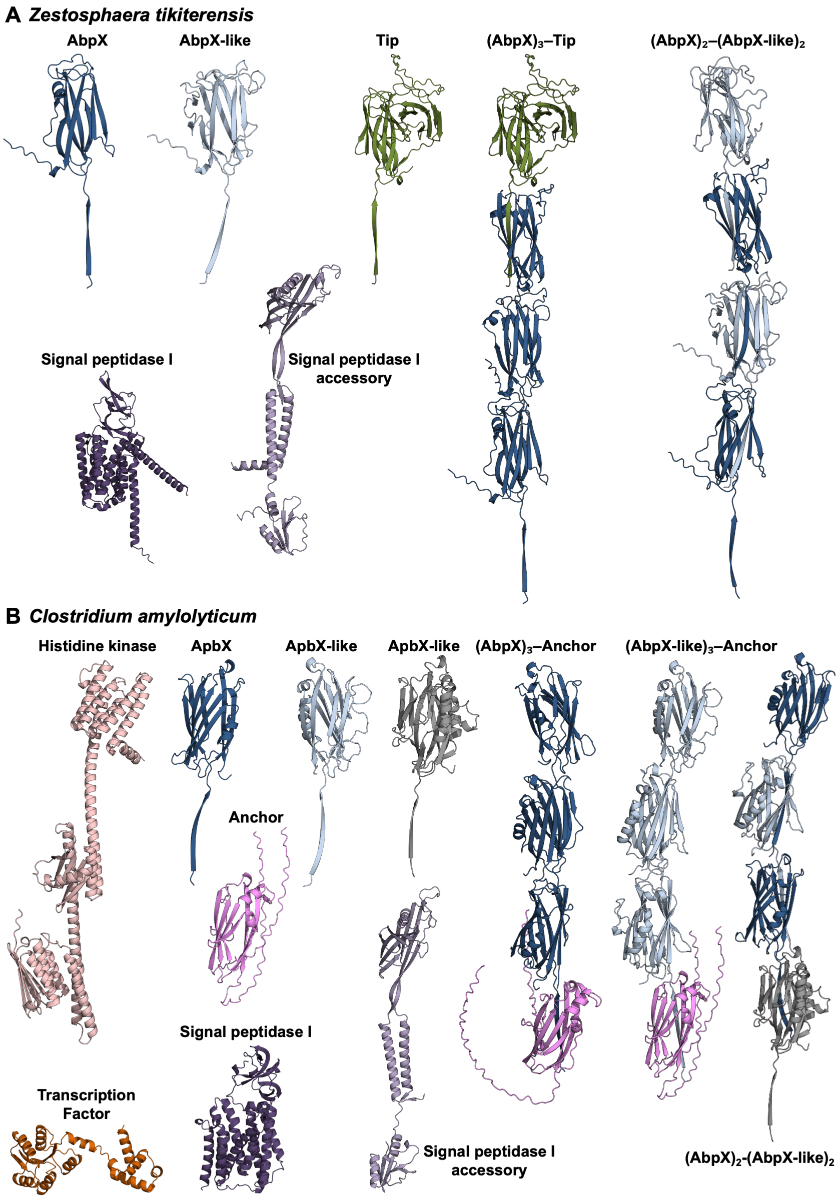


**Fig. S16.** (**A**) Predicted structures for the *Zestosphaera tikiterensis abpX* genomic locus, including AbpX (UniProt: A0A2R7Y7S4), the AbpX-like minor filament component (A0A2R7Y7T0), the tip protein (A0A2R7Y9R0), SPI (A0A2R7Y7S8), and the SPI-accessory protein (A0A2R7Y7T3). Monomeric models are shown for all components, while predicted multimeric assemblies are provided only for AbpX, the AbpX-like minor filament component, and the tip protein. (**B**) Predicted structures for the *Clostridium amylolyticum abpX* genomic locus, including the LPXTG-type cell-wall anchor (A0A1M6GRE1), the histidine kinase (A0A1M6GRG9), a Spo0A-family transcription factor (A0A1H9ZYW4), SPI (A0A1M6GRN4), AbpX (A0A1M6GRR4), two AbpX-like minor filament components (A0A1M6GRY3, A0A1M6GRP1), and the SPI-accessory protein (UniProt: A0A1M6GRS4). Monomeric models are shown for all components, while predicted multimeric assemblies are provided only for AbpX, the AbpX-like minor filament components, and the LPXTG-type anchor.

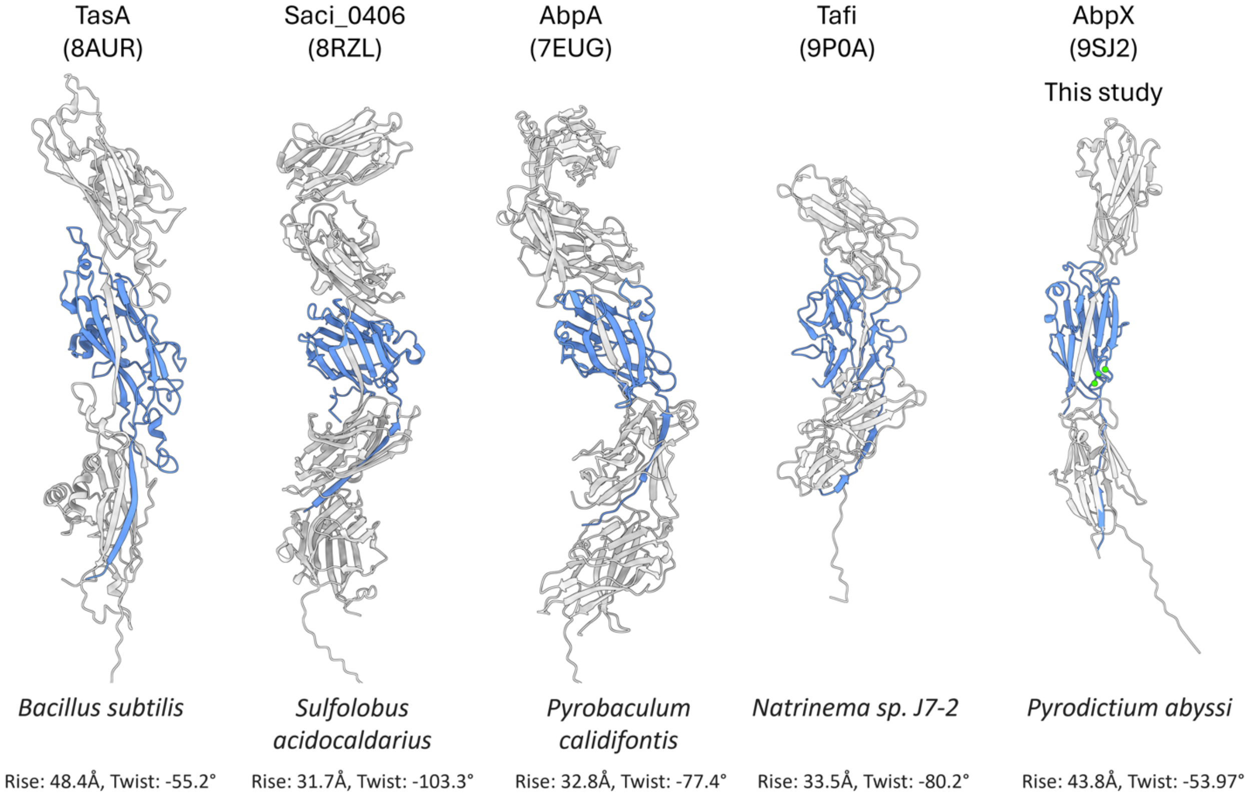


**Fig. S17. The structural repertoire of donor-strand complemented protein polymers derived from archaeal and bacterial bundling pili.**

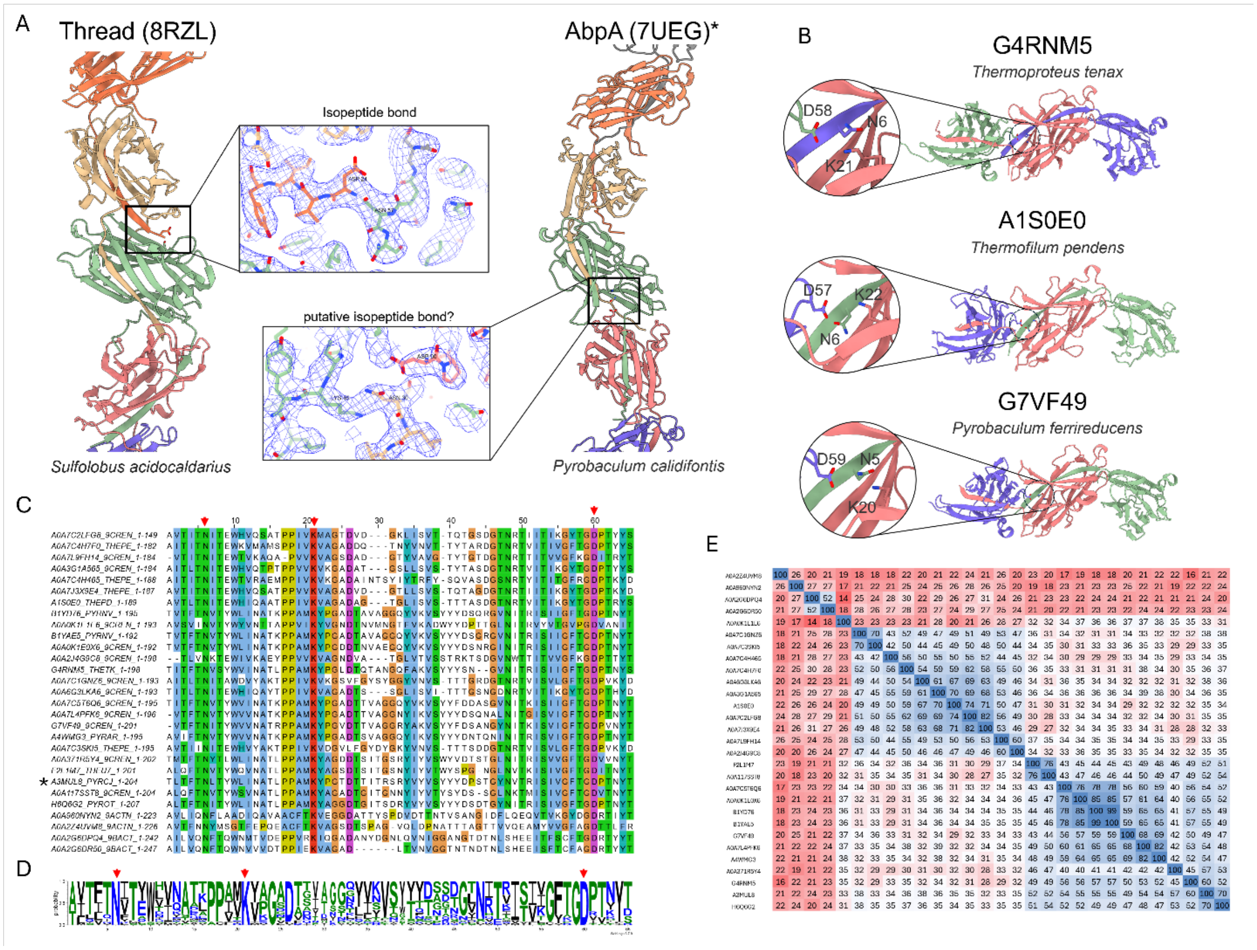


**Fig. S18 Inter-subunit isopeptide bonds in archaeal donor-strand complemented filaments.** (**A**) The thread filament of *Sulfolobus acidocaldarius* (PDB: 8RZL) is stabilized by an isopeptide bond between the N-terminal ASP24 and ASN57. The archaeal bundling pilus AbpA of *Pyrobaculum calidifontis* (PDB: 7UEG) has no reported isopeptide bonds, but we identified clear connected density in the cryoEM reconstruction EMD-26474, between residues ASN30 and LYS45, with ASP90 in proximity. Given the limited resolution of EMD-26474 (4.0 Å), it is impossible to draw firm conclusions regarding this putative IPB site. (**B**) AlphaFold3 predictions of 3 selected homologues of AbpA demonstrating structural conservation of the N/K/D triad. (**C**) Multiple sequence alignment of AbpA homologues found with blastp demonstrates strict sequence conservation of the N, K, D triad. AbpA is marked with an asterisk; (**D**) Corresponding weblogo of the MSA in (**C**). (**E**) Pairwise sequence identity matrix of the sequences in (**C**) highlighting low sequence identity across the AbpA homologues.

**Table S1. CryoEM data collection and refinement statistics for AbpX fibrils.**

| Parameter | AbpX |
|---|---|
| Voltage (kV) | 300 |
| Electron exposure ($e^{-}$ Å$^{-2}$) | 60 |
| Pixel size (Å) | 0.695 |
| Particle images (n) | 1,572,118 |
| Shift (pixel) | 20 |
| Point group | C1 |
| Helical rise (Å) | 44.01 |
| Helical twist (°) | -53.97 |
| Map:map FSC (0.143) | 3.43 |
| Model:map FSC (0.50) | 3.94 |
| $d_{99}$ | 4.20 |
| Ramachandran Favored (%) | 98.74 |
| Ramachandran Outliers (%) | 0.0 |
| RSCC | 0.82 |
| Clash score | 7.39 |
| Bonds RMSD, length (Å) | 0.003 |
| Bonds RMSD, angles (°) | 0.631 |
| **Deposition ID** | |
| PDB (model) | 9SJ2 |
| EMDB (map) | EMD-54935 |

**Table S2. Crystallographic data collection and refinement statistics for sc-AbpX.** Values in parenthesis refer to the highest recorded resolution shell.

| **Data collection** | | **Refinement** | |
|---|---|---|---|
| Space group | P 65 2 2 | Resolution (Å) | 1.49 |
| Cell dimensions | | # unique reflections | 16094 (145) |
| a, b, c (Å) | 55.45 55.45 268.59 | R-work/R-free | 0.203/0.249 |
| α, β, γ (°) | 90 90 120 | Number of atoms | 1310 |
| Resolution (Å)* | 48.024 - 1.491 | protein | 1215 |
| $R_{pim}$ | 0.015 (0.307) | ligands/ions | 3 |
| $R_{merge}$ | 0.090(1.367) | water | 92 |
| I/σ(I) | 18.6 (2.1) | Average B-factors | 35.97 |
| Completeness (%) | 91.4 (69.5) | protein | 35.39 |
| Multiplicity | 34.2 (19.1) | ligands/ions | 49.84 |
| | | water | 43.09 |
| | | RMS deviations | |
| | | Bond angles (°) | 0.545 |
| | | Bond length (Å) | 0.003 |
| | | Ramachandran | |
| | | favored (%) | 97.59% |
| | | allowed (%) | 2.41% |
| | | outliers (%) | 0 % |
| | | **PDB code** | 9RHP |

**Table S3. Accession IDs for proteins encoded by archaeal and bacterial *abpX* loci.**

| Organism | Protein IDs |
| --- | --- |
| *Pyrodictium abyssi* | WP_338249486.1, WP_338249487.1 |
| *Methanosuratincola subterraneus* | A0A444L6T6, A0A3S3SRM5, A0A3S3RZT5, A0A3S3TRS3, A0A3S3RBX0, A0A3S3S7L5 |
| *Zestosphaera tikiterensis* | A0A2R7Y7S4, A0A2R7Y7T0, A0A2R7Y9R0, A0A2R7Y7S8, A0A2R7Y7T3 |
| *Ignisphaera aggregans* | E0SRZ6, E0SRZ5, E0SRZ4, E0SRZ3, E0SRZ2 |
| *Pyrobaculum ferrireducens* | G7VCD2, G7VCD3, G7VCD4, G7VCD5, G7VCD6 |
| *Desulfurococcus amylolyticus* | B8D534, B8D533, B8D532, B8D531, B8D530, B8D529 |
| *Ammonifex thiophilus* | A0A3D8P4F6, A0A3D8P4D9, A0A3D8P2P2, A0A3D8P4Q7, A0A3D8P2M4, A0A3D8P2J4 |
| *Clostridium amylolyticum* | A0A1M6GRE1, A0A1M6GRG9, A0A1M6GRN4, A0A1M6GRR4, A0A1M6GRY3, A0A1M6GRP1, A0A1M6GRS4 |
| *Natronincola peptidivorans* | A0A1H9ZYW4, A0A1H9ZYU5, A0A1I0A021, A0A1I0A1I3, A0A1H9ZYY9, A0A1H9ZZ32, A0A1H9ZZL4, A0A1H9ZZB4, A0A1I0A0H4 |
| *Thermacetogenium phaeum* | K4LH80, K4LK71, K4LWH1, K4LHJ6, K4LJX3, K4LH75, K4LK66 |
| *Syntrophothermus lipocalidus* | D7CMU8, D7CMU7, D7CMU6, D7CMU5, D7CMU4, D7CMU3, D7CMU2, D7CMU1, D7CMU0, D7CMT9 |
| *Heliobacterium mobile* | A0A6I3SMJ2, A0A6I3SP27, A0A6I3SM01, A0A6I3SM80, A0A6I3SM17, A0A6I3SM16, A0A6I3SN78, A0A6I3SM09, A0A6I3SM31, A0A6I3SMP6 |

**Movie S1.**

**Structural conservation of para-crystalline AbpX fiber bundles.** CryoEM 2D class averages of para-crystalline fibril bundles of recombinant AbpX, with the corresponding power spectrum of the 2D class average. The position of the first layer line (1/271Å) and the n = 0 maximum on the meridional (1/44Å) indicates structural consistency at the level of the helical rise and pitch of the protofilaments that are embedded within the para-crystalline bundles.

**Movie S2.**

**Axial flexing of the AbpX filament.** 3D representation of the result of cryoSPARCs “3D variability analysis” using the 1,572,118 cryoEM particles that were used for the helical reconstruction of AbpX. The search was conducted for 3 principal modes (0,1,2), while limiting the analysis to 5 Å resolution. The video displays a series of 20 reconstructions that were generated along mode 2, using the cryoSPARC job “3D variability display” set to “simple” mode. The 20 volumes were loaded as a volume series into ChimeraX 1.9 and exported into mp4 format using the “movie” command.

**Data S1.**

**Amino acid sequences of proteins encoded by archaeal and bacterial *abpX* loci listed in Table S3.**
